# Genome-wide analysis of consistently RNA edited sites in human blood reveals interactions with mRNA processing genes and suggests correlations with cell types and biological variables

**DOI:** 10.1101/254045

**Authors:** Edoardo Giacopuzzi, Massimo Gennarelli, Chiara Sacco, Alice Filippini, Jessica Mingardi, Chiara Magri, Alessandro Barbon

## Abstract

**Background:** A-to-I RNA editing is a co-/post-transcriptional modification catalyzed by ADAR enzymes, that deaminates Adenosines (A) into Inosines (I). Most of known editing events are located within inverted ALU repeats, but they also occur in coding sequences and may alter the function of encoded proteins. RNA editing contributes to generate transcriptomic diversity and it is found altered in cancer, autoimmune and neurological disorders. Emerging evidences indicate that editing process could be influenced by genetic variations, biological and environmental variables.

**Results:** We analyzed RNA editing levels in human blood using RNA-seq data from 459 healthy individuals and identified 2,079 sites consistently edited in this tissue. As expected, analysis of gene expression revealed that *ADAR* is the major contributor to editing on these sites, explaining ∼13% of observed variability. After removing *ADAR* effect, we found significant associations for 1,122 genes, mainly involved in RNA processing. These genes were significantly enriched in genes encoding proteins interacting with ADARs, including 276 potential ADARs interactors and 9 ADARs direct partners. In addition, our analysis revealed several factors potentially influencing RNA editing in blood, including cell composition, age, Body Mass Index, smoke and alcohol consumption. Finally, we identified genetic loci associated with editing levels, including known *ADAR* eQTLs and a small region on chromosome 7, containing *LOC730338,* a lincRNA gene that appears to modulate ADARs mRNA expression.

**Conclusions:** Our data provides a detailed picture of the most relevant RNA editing events and their variability in human blood, giving interesting insights on potential mechanisms behind this post-transcriptional modification and its regulation in this tissue.

## Background

RNA editing is a co-/post-transcriptional process based on the enzymatic deamination of specific adenosines (A) into inosines (I). Since inosine has similar base-pairing properties to guanosine, it is read as guanosine by both splicing and translation machineries, thus generating different RNA molecules from those coded by DNA [1]. RNA editing contributes to the diversification of the information that is encoded in the genome of an organism, thereby providing a greater degree of complexity. Currently, the conversion of A to I is thought to be the most common RNA editing process in higher eukaryotic cells [2].

RNA editing is catalyzed by adenosine deaminase enzymes (ADARs) [3, 4]. In mammals, three members of the ADAR family have been characterized so far. ADAR1 (gene name: *ADAR*) and ADAR2 (gene name: *ADARB1*) are active enzymes expressed in many tissues, while ADAR3 (gene name: *ADARB2*) is expressed specifically in the Central Nervous System (CNS). To date, no functional RNA editing activity has been attributed to this enzyme. The critical role of ADAR enzymes is shown by phenotypes of knockout mice that resulted in embryonic lethality or death shortly after birth [5–7], clearly indicating that A-I RNA editing is essential for normal life and development. In addition, dysregulated RNA editing levels at specific re-coding sites have been linked with a variety of diseases, including neurological or psychiatric disorders and cancer [2, 8, 9]. Interestingly, ADARs mRNA and protein expression levels do not always reflect RNA editing levels [10]. It has been shown that the subcellular distribution of ADAR enzymes [11] and their interaction with inhibitors [12, 13] and activators [14, 15] influence ADARs activities.

Originally, A-to-I RNA editing in mammalian cells was described for a low number of mRNAs and it was responsible for deep changes of protein functions. These editing sites were discovered serendipitously by directly comparing DNA and cDNA sequences [16, 17]. The number of identified RNA editing sites has largely increased with the widespread adoption of RNA sequencing (RNA-seq), reaching over two million sites. The majority of RNA editing sites is located within intragenic non-coding sequences: 5’UTRs, 3’UTRs and intronic retrotransposon elements, such as ALU inverted repeats [18, 19]. With lowering cost of NGS, many RNA-Seq datasets from human tissues, healthy and pathological conditions, have been deposited in sequence databases, available to the scientific community. In parallel, the development of computational pipelines to search for RNA editing sites on RNA-Seq data, allowed a global analysis of the editing reaction, shedding light on its evolutionary conservation [20], tissues specificity [21, 22], cellular specificity [23] and its role in diseases such cancer [24] or neurological disorders [9, 25].

About 2.5 million editing sites have been identified so far and are listed in RNA editing databases [26, 27], but only recently the dynamic and regulation of RNA editing has been systematically investigated in human tissues [22]. However, little is known about how editing process could be influenced by genetic variations [28, 29], biological and environmental variables [30]. Here, we want to go further in characterizing and understanding the complexity of RNA editing. Focusing on the most likely biologically relevant sites, we sought to unveil possible correlations with gene expression and genetic variations. To this aim, we investigated consistently edited sites from existing RNA seq dataset of whole blood from 459 healthy subjects [31], correlating editing levels with blood cellular composition, with a collection of 28 biological and pharmacological variables, as well as with genes expression and genotyping data.

## Results

### RNA editing sites consistently edited in human blood samples

Of the ∼ 2M editing sites reported in RADAR database, 709,184 sites have an adequate coverage (> 10 reads) in our dataset of 459 RNA-seq, covering > 75% of the total sites reported for genes expressed in blood according to GTEx data (Additional file 1: Figure S1). Most of these sites are edited only in a small fraction of samples and 691,304 (97.5%) have no detectable editing levels in our cohort (Additional file 1: Figure S2). To provide a picture of the most biologically relevant editing sites in human blood we focused our attention on 2,079 consistently editing sites (CES), namely those with at least 5% of editing level in at least 20% of individuals. These sites are distributed across 421 genes and mainly localized in ALU regions (1,805; 86.5%) and 3’UTR regions (1,234; 59.4%). Overall, we detected 1,266 sites in exons of protein coding genes, including 10 recoding sites (resulting in a missense substitution) and 12 synonymous sites. We also detected 53 sites annotated on ncRNAs (Figure 1a, b). Detailed statistics of the 2,079 trusted sites are reported in Additional file 2, while recoding sites are reported in Table 1.

**Table 1.** Editing levels detected for the 10 recoding sites identified in human blood

| Site id (hg19) | Gene | Strand | Aa change | Alu | Editing Minimum | Editing Mean | Editing Maximum |
| --- | --- | --- | --- | --- | --- | --- | --- |
| chr3_49398423 | *RHOA* | - | Lys->Arg | yes | 0.19 | 0.33 | 0.64 |
| chr4_2835556 | *SH3BP2* | + | Arg->Gly | no | 0.05 | 0.08 | 0.16 |
| chr4_2940026 | *NOP14* | - | Asn->Ser | yes | 0.1 | 0.23 | 0.56 |
| chr4_77979680 | *CCNI* | - | Arg->Gly | no | 0.05 | 0.08 | 0.19 |
| chr8_103841636 | *AZIN1* | - | Ser->Gly | no | 0.07 | 0.16 | 0.45 |
| chr13_52604264 | *UTP14C* | + | Ser->Gly | no | 0.24 | 0.64 | 1 |
| chr13_52604880 | *UTP14C* | + | Gln->Arg | no | 0.45 | 0.85 | 1 |
| chr15_75646086 | *NEIL1* | + | Lys->Arg | no | 0.27 | 0.73 | 1 |
| chr16_3292200 | *MEFV* | - | Stp->Trp | yes | 0.05 | 0.16 | 0.36 |
| chr20_36147563 | *BLCAP* | - | Gln->Arg | no | 0.05 | 0.14 | 0.33 |

**Figure 1.**
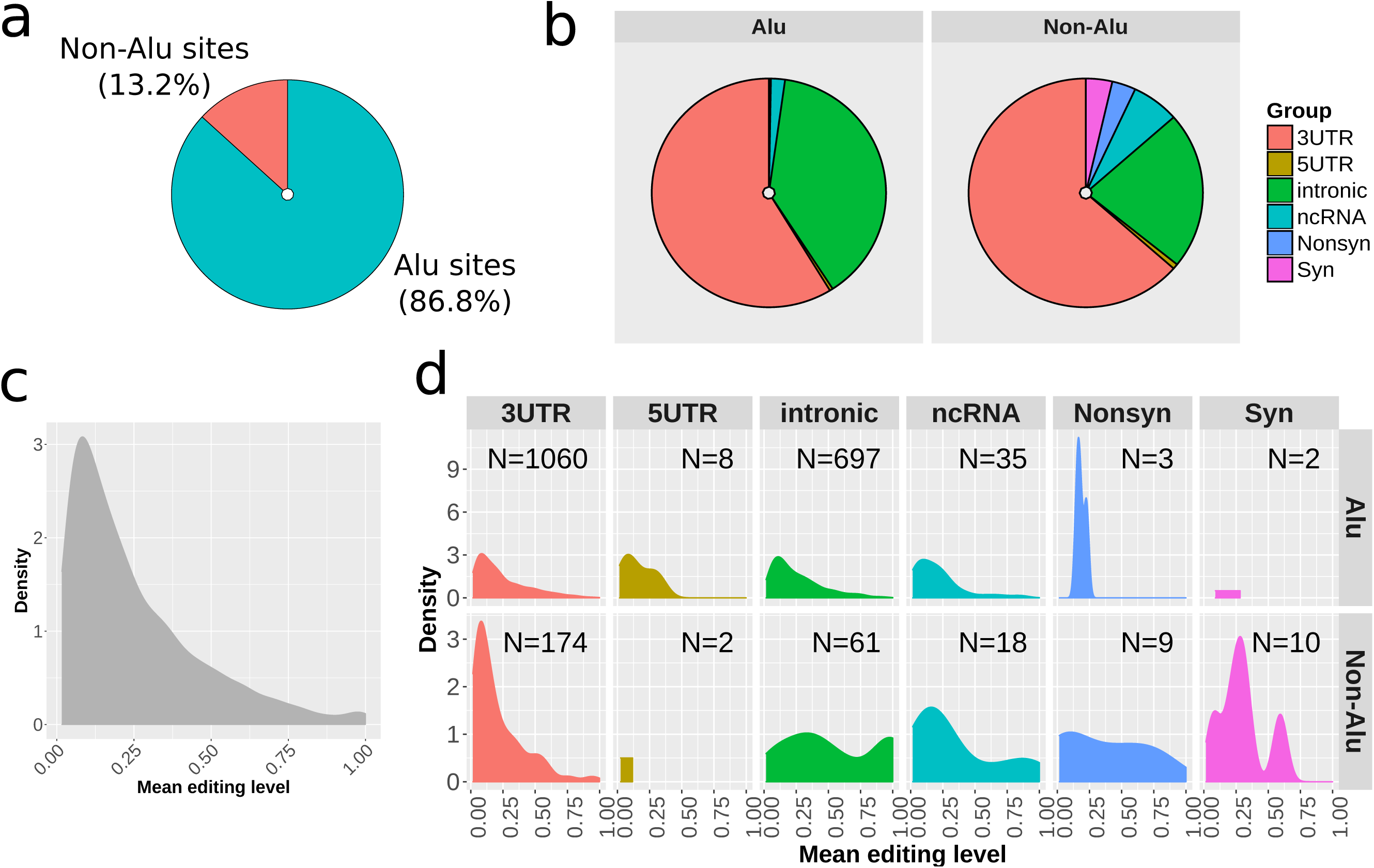
Distribution of 2,079 consistently edited sites (CES) analyzed in the study (a) Distribution of the 2,079 CES within ALU regions and (b) based on functional classification. (c) Density plot representing overall distribution of editing levels. (d) Density plots of editing levels for different editing site categories and ALU/non-ALU sites.

Considering mean values for each site, detected editing levels range from 0.05 to 1, with most sites showing moderate editing levels between 0.05 and 0.30 (Figure 1c). We also detected 33 sites highly edited (mean value ≥ 0.9), located mainly in intronic regions (Figure 1d). Highly edited sites are reported in Additional file 1: Table S1. To further assess reliability of detected sites, we compared the CES editing levels with those reported in the REDIportal database [26], a well-established resource containing multi-tissue estimations of RNA editing levels. When considering the REDIportal blood tissue data, the comparison revealed high concordance (concordance correlation coefficient 0.84, Additional file 1: Figure S3) for 2,003 overlapping sites, with 20 out of 33 highly edited sites (60 %) showing similar editing levels. When we excluded sites measured only in a single subject in REDIportal, we found that 16 out of 18 sites (89 %) have high level of editing in both datasets, suggesting that our highly edited sites are probably true editing events rather than systematic sequencing errors. However, the occurrence of sequencing artifact could not be completely excluded. For the sites included in this study, the editing levels from REDIportal are reported in Additional file 2.

We found 495 CES located within known miRNA binding sites from TargetScan v.7.2. Among these, 466 (94%) are located in 3’ UTR (representing 37.7% of the total 3’UTR sites); however only 4 CES sites overlap with conserved binding sites for broadly conserved miRNAs (Additional file 1: Figure S4). Broadly conserved miRNAs are defined as conserved across most vertebrates, usually to zebrafish, while binding site conservation is defined by conserved branch length, with each site type having a different threshold for conservation. The overlap with miRNA binding sites is reported for each editing site in Additional file 2.

We used Spearman correlation test to analyze correlation in editing level changes across the 2,079 CES to find sites with co-regulated RNA editing. We found 270 significant relationships (FDR < 0.05) involving 361 sites. Correlations were generally low with only 58 sites with relationships above 0.5 rho value. Correlations become stronger for close sites, especially below 50 bp distance, with 30 out of 33 (91%) high rho (≥ 0.5) relationships located in this range. Considering the 100 middle level correlations (rho between 0.3 and 0.5), 95 are observed between sites within about 1 kb distance and 83 between sites in the 50 bp range. Interestingly, we also observed 5 relationships between sites on different chromosomes. No strong negative correlations (rho < −0.5) were observed (Additional file 1: Figure S5). Full results of correlation analysis are report in Additional file 3.

### Genes influencing the total editing rate of CES

We performed regression analysis to identify genes whose expression is associated to the CES total editing rate, calculated for each subject as the total sum of G-containing reads divided by the total number of reads observed at all the 2,079 CES. To avoid biases due to the influence of different blood cell type compositions, we used normalized gene expression values provided in [31], where the effect of cell type composition was regressed out from read counts using ridge regression. The analysis revealed 4,719 genes associated with the CES total editing rate (FDR < 0.05). Enrichment analysis on Gene Ontology biological processes (GO-BP) revealed a strong enrichment for genes involved in immune system and interferon signaling (FDR < 1e-6, Figure 2a). Among significant genes, *ADAR* emerged as the top influencing factor, explaining about 13% of the observed variability, while *ADARB1* showed no significant effect (Figure 2b). The influence of *ADAR* was similar on ALU (∼10%) and non-ALU (∼13%) sites, while *ADARB1* remains not associated when considering the two groups separately (Additional file 1: Figure S6). *ADARB2* gene was not detectable in our gene expression data. When the same analysis was repeated removing *ADAR* effect, we obtained 1,122 genes associated with CES total editing rate (FDR < 0.05), including 376 with a strong association at FDR < 0.01 (Additional file 4). Enrichment analysis on GO-BP and REACTOME pathways revealed that these genes mainly impact ribonucleoprotein complex biogenesis and RNA metabolism / processing (Figure 2c).

**Figure 2.**
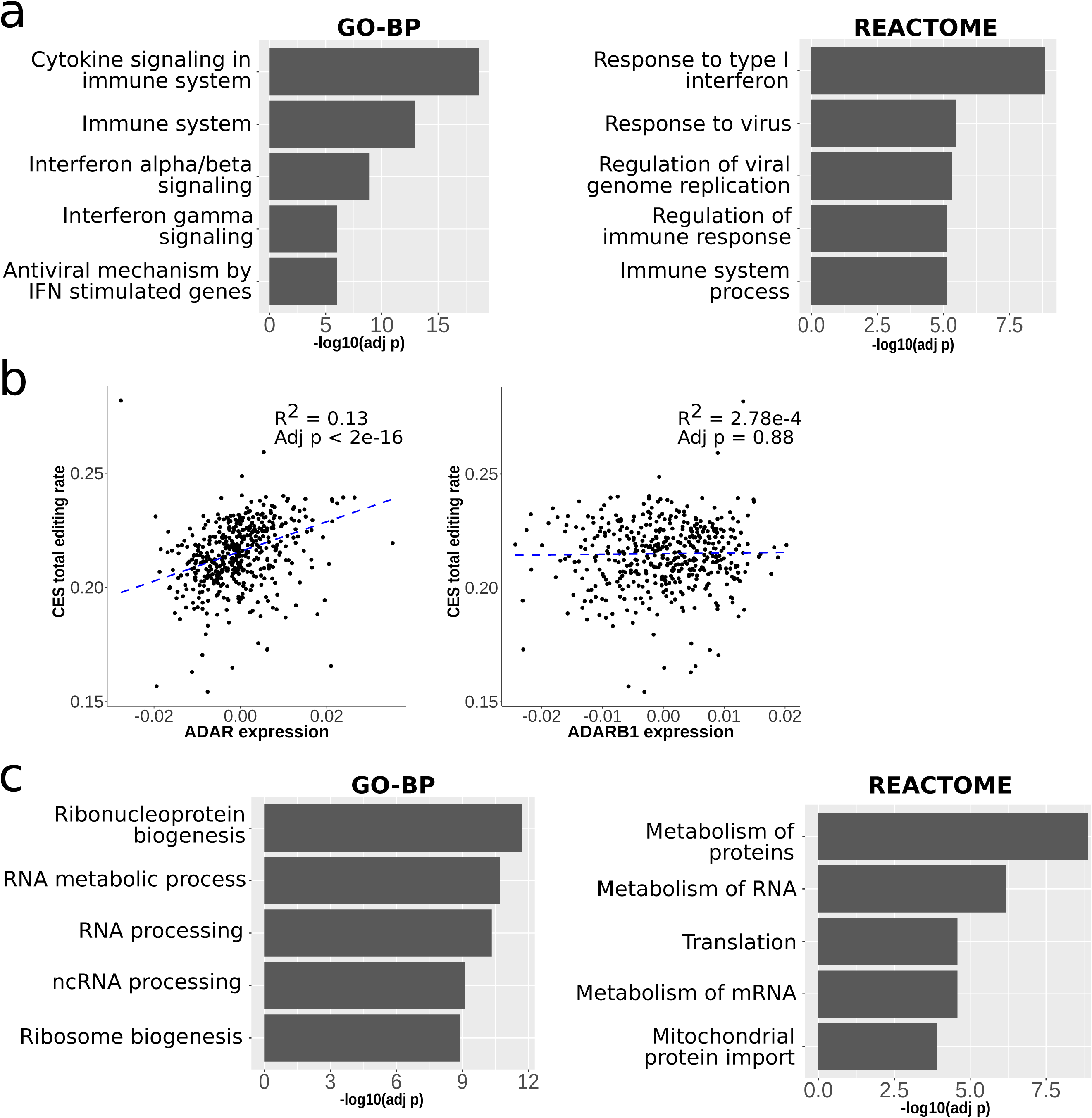
Association between gene expression and CES total editing rate We analyzed association between CES total editing rate and gene expression for 14,961 human genes. (a) Gene set enrichment analysis by hypergeometric test on GO-BP categories and REACTOME pathways revealed that associated genes are mainly involved in immune system response mediated by interferon I and alpha / beta. (b) When we analyze distribution of CES total editing rate and *ADAR* gene expression, *ADAR* expression levels explains ∼ 13% of observed variability. No significant effect is observed for *ADARB1* expression. *ADAR* and *ADARB1* expression levels are reported as residuals of ridge regression with technical covariates (see description of data in methods section). The graphs report adjusted p-value and R2 value from robust regression analysis. (c) The 1,122 genes associated to CES total editing rate after removing *ADAR* expression effect were enriched for genes mainly involved in ribonucleoprotein and RNA processing.

To assess possible interactions between ADAR enzymes and genes whose expression is associated with CES total editing rate, we performed network analysis using data on protein-protein interactions from STRING v.10, BioPlex and BIOGRID databases. Among the proteins encoded by the 376 genes significantly associated with CES total editing rate (FDR < 0.01), 285 (76 %) were connected to ADAR1 or ADAR2, directly or through a first-level interactor. The resulting network includes a total of 415 proteins: 285 encoded by genes significantly associated with CES, 2 ADARs proteins and 128 added partners (first-level interactors which connect significant proteins to ADARs, but are not encoded by significant genes) (Figure 3a). The observed fraction of ADARs-connected proteins (285 out of 376) represents a significant enrichment compared to random groups (empirical p-value < 1e-06, Figure 3b) and these proteins are strongly enriched for RNA binding proteins (Figure 3c). Among the 285 genes significantly associated to CES total editing rate, we identified 9 genes encoding proteins with direct interactions with ADARs (Table 2). We estimated the role of each node in this network looking at degree and betweenness values. Degree value accounts for the number of interactions (edges) involving a single node in the network, while betweenness is a measure of centrality based on shortest paths. Nodes with high values of betweenness centrality would have a more relevant role in the network since an increased proportion of the connections between distant nodes passes through them. Among ADARs proteins, ADAR1 has considerably more network interactions (0.077 betweenness centrality, 72 degree values) compared to ADAR2 (0.018 betweenness centrality, 29 degree). Among genes associated with CES total editing rate, those encoding for proteins with a direct interaction with ADARs, *ELAVL1, RPA1* and *IFI16* act as relevant hub nodes, with betweenness centrality values of 0.137, 0.028, 0.020, respectively (Figure 3d). Detailed network-based statistics are reported in Additional file 5, together with adjusted p values for association with CES total editing rate. Since a direct interaction of ADAR1 with IFI16 and RPA70 (encoded by *RPA1*) proteins has never been reported, we decided to experimentally verify these results by co-immunoprecipitation experiments in Epstein-Barr Virus (EBV)-immortalized human B cell lines (B-EBV) (Figure 3e). The results confirmed that IFI16 protein is an interactor of ADAR1, at least at low level, as indicated by the clear band of interaction. Considering RPA70, the protein appeared in ADAR1 precipitate, but a faint band with a similar molecular weight is also present in IgG precipitate, suggesting that these results may need further investigation.

**Figure 3.**
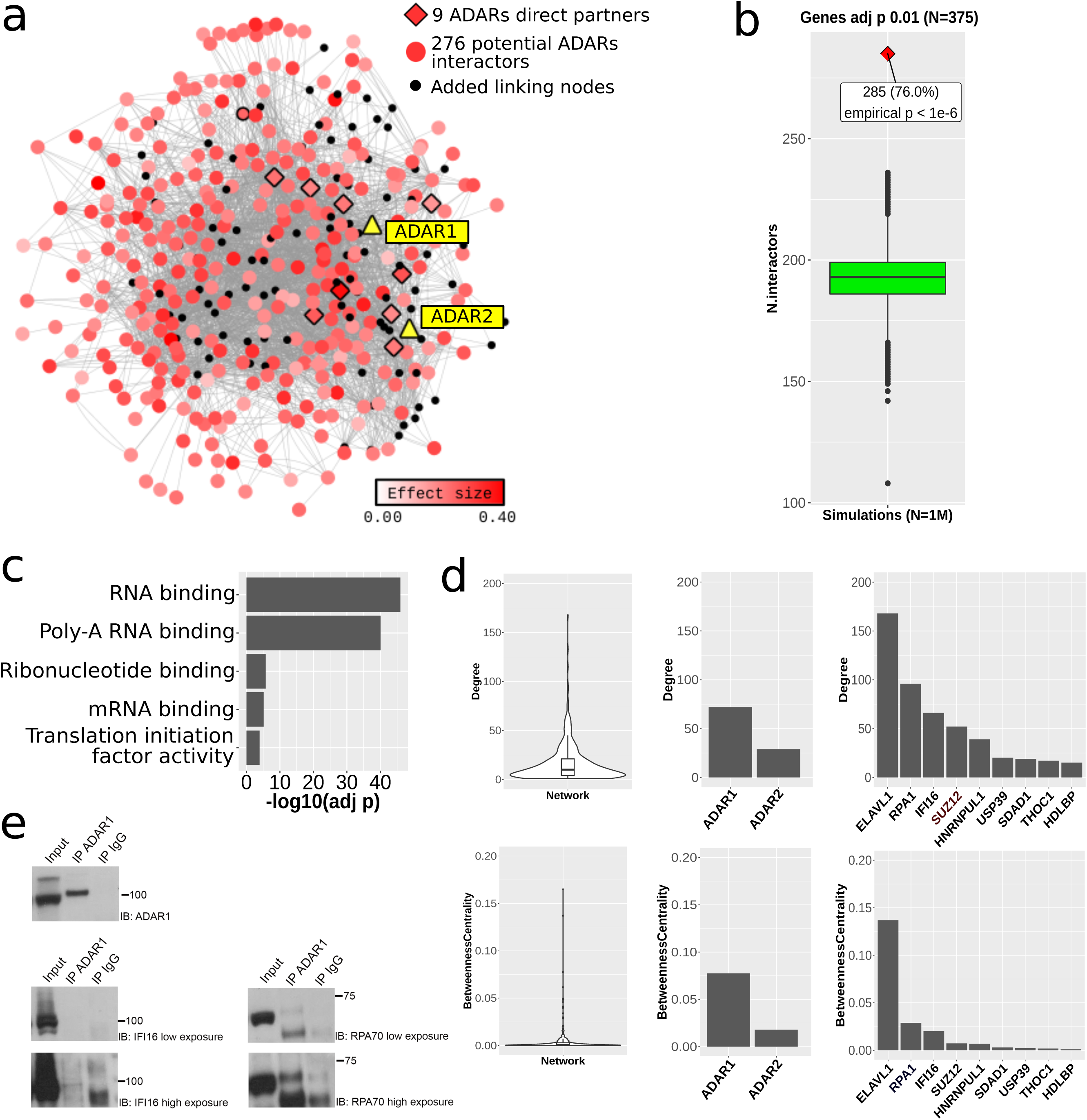
Genes associated with CES total editing rate are enriched for ADAR interactors (a) Reconstructed PPI network including ADARs and proteins encoded by best genes significantly associated with global editing levels (FDR < 0.01). Among these proteins, we observed 285 potential ADARs interactors, including 9 direct partners of ADARs proteins. (b) Boxplot of number of ADARs interacting genes observed in 1M random simulations. The observed number of interactions (285) resulted in empirical p-value < 1e-6. (c) ADARs interactors are strongly enriched for RNA binding proteins in GO-MF categories. (d) Distribution of degree and betweenness centrality values among network nodes are represented by violin plots. ADAR1 protein has a major role (higher values) among ADAR proteins. Among ADARs direct partners, *ELAVL1, RPA1* and *IFI16* showed high values of degree and betweenness centrality, suggesting a central role in the network. (e) ADAR1 interaction with RPA70 (coded by RPA1) and IFI16 determined by co-immunoprecipitation. After immunoprecipitation with ADAR1 antibody, western blot for IFI16 and RPA70 are reported. For a better discrimination two times of exposure are reported in the figure.

**Table 2.**
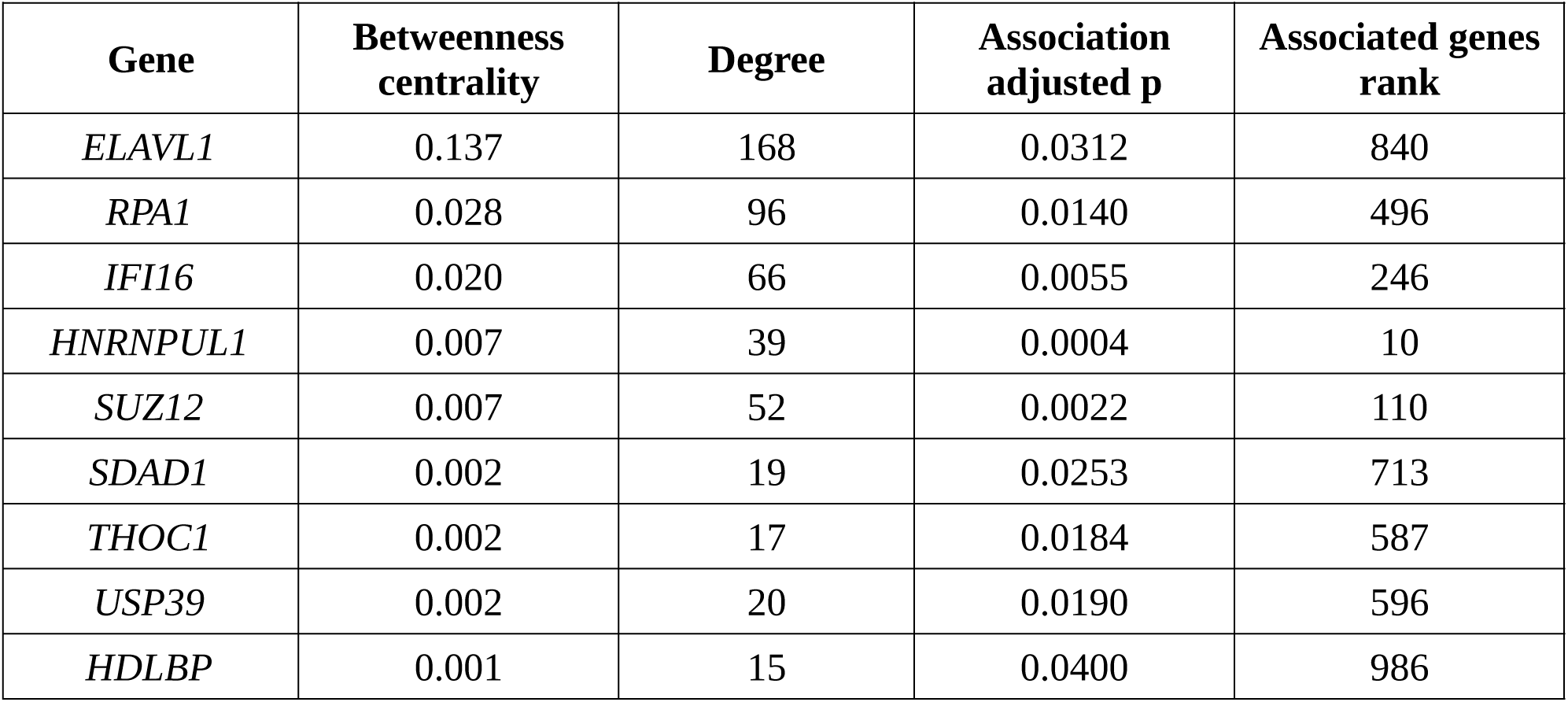
Network based statistics for the 9 ADARs direct partners significantly associated to global editing levels (adjusted p < 0.05)

| Gene | Betweenness centrality | Degree | Association adjusted p | Associated genes rank |
| --- | --- | --- | --- | --- |
| *ELAVL1* | 0.137 | 168 | 0.0312 | 840 |
| *RPA1* | 0.028 | 96 | 0.0140 | 496 |
| *IFI16* | 0.020 | 66 | 0.0055 | 246 |
| *HNRNPUL1* | 0.007 | 39 | 0.0004 | 10 |
| *SUZ12* | 0.007 | 52 | 0.0022 | 110 |
| *SDAD1* | 0.002 | 19 | 0.0253 | 713 |
| *THOC1* | 0.002 | 17 | 0.0184 | 587 |
| *USP39* | 0.002 | 20 | 0.0190 | 596 |
| *HDLBP* | 0.001 | 15 | 0.0400 | 986 |

### Influence of cell composition on the total editing rate of CES

To assess how changes in cell composition of whole blood can affect the observed editing levels, we correlated the proportions of different blood cell types (as provided in [31]) with the total editing rate of CES. Among the 7 cell types considered, 4 showed a significant correlation with total editing rate, namely T helper (Th), monocytes, dendritic cells (DC) and neutrophils (Figure 4). We also found 4 cell types associated with *ADAR* expression (neutrophils, Th, Natural Killer and B) and 4 with *ADARB1* expression (neutrophils, monocytes, Th and B). Intriguingly, the associations of *ADAR* and *ADARB1* with cell types are always in opposite directions. See Additional File 1: Table S2 for complete results. Overall, cell composition variables explain 19% and 53% of *ADAR* and *ADARB1* expression variability, respectively (Additional file 1: Figure S7)

**Figure 4.**
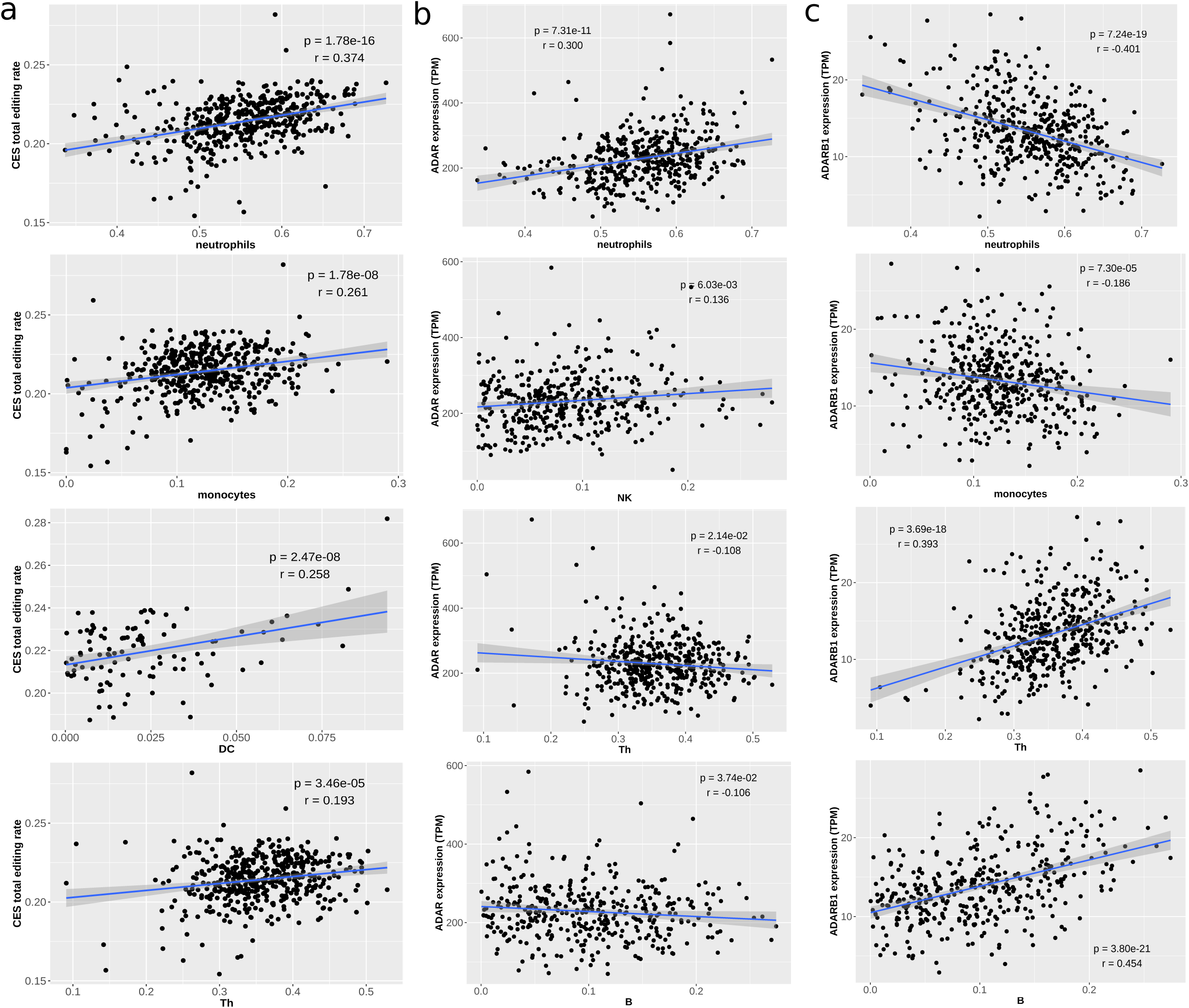
Impact of cell composition on CES total editing rate and *ADAR / ADARB1* expression Our analysis revealed strong associations with CES total editing rate for 4 cell type variables (a), representing proportion of neutrophils, monocytes, dendritic cells (DC) and T helper (Th). Specific cell variables resulted significantly associated also to *ADAR* (b) and *ADARB1* (c) expression. Significance level (p) and correlation coefficient (r) are reported in each plot based on Pearson’s product-moment. Only non-zero observations are plotted.

### Biological factors possibly influencing editing levels

In order to identify possible biological factors influencing editing levels, we studied the correlation of the 28 biological / pharmacological variables described in Additional file 1: Table S3 with CES total editing rate and with the *ADAR* expression level. Overall, 5 variables revealed a significant correlation with CES total editing rate and 3 of them remained associated even after correction for cell type composition, namely blood pressure medications, age and current Body Mass Index (BMI) (Figure 5). We also found 5 variables significantly correlated with the expression level of either *ADAR* (blood pressure medications, current and max BMI, age and sex) or *ADARB1* (sex, time of draw, thyroid medications, ate before and proton-pump inhibitors), even if the effect was generally small after correction for cell composition (Additional File 1: Table S4 and Figure S7).

**Figure 5.**
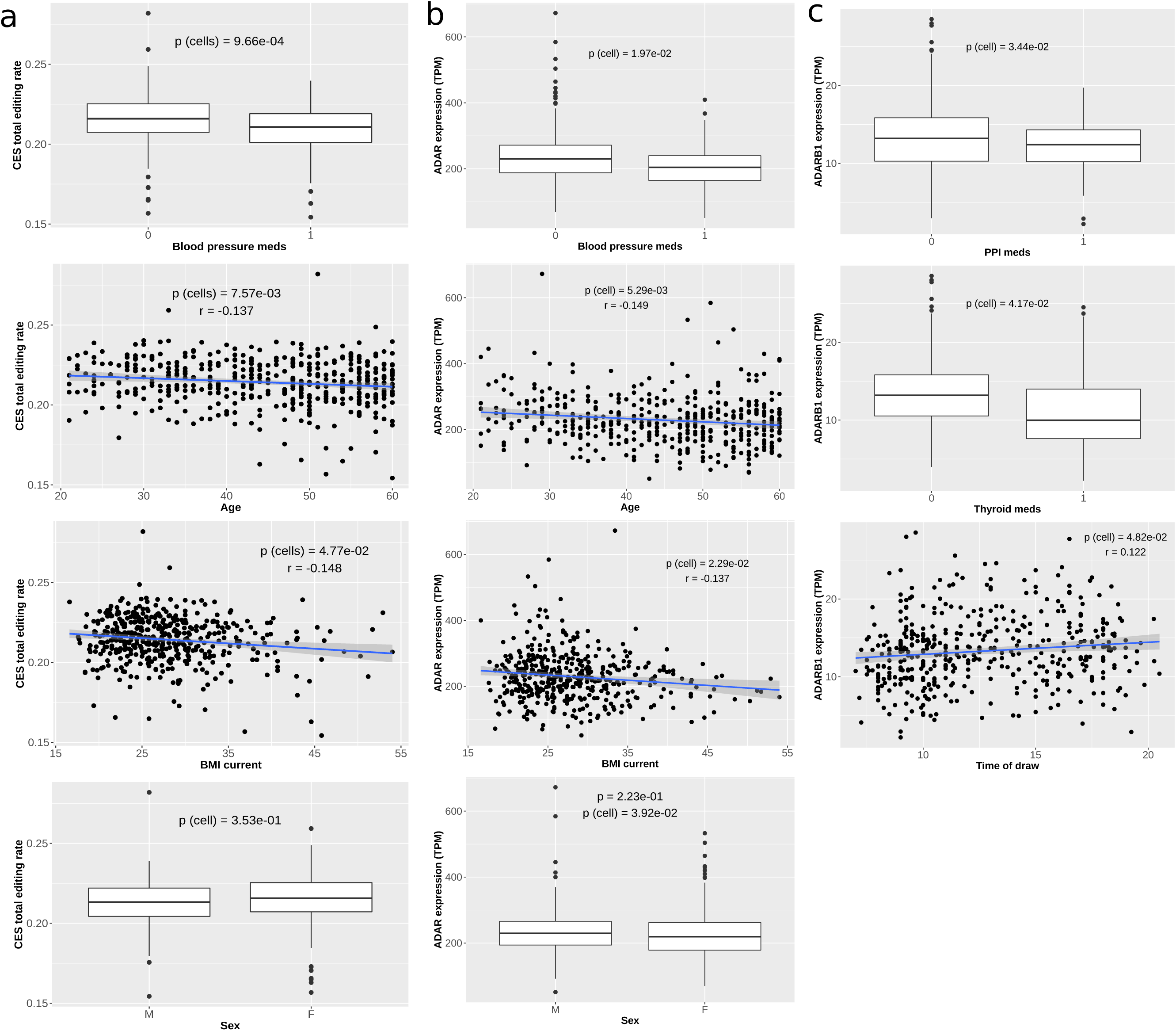
Impact of biological / pharmacological factors on CES total editing rate and *ADAR / ADARB1* expression Our analysis revealed significant associations with CES total editing rate for blood pressure medication, BMI current, Age and Sex (a). Specific biological variables resulted significantly associated also to *ADAR* (b) and *ADARB1* (c) expression. Significance level of association after correction for cell composition is reported (p (cell)) is reported in each plot based on Mann-Whitney-Wilcoxon or Pearson’s product-moment correlation test for binary and continuous variables, respectively. For continuous variables the Pearson correlation coefficient (r) is also reported.

### Principal component analysis

To better investigate the effect of cell type composition, biological / pharmacological variables and *ADAR* expression and identify correlations between these variables and specific groups of CES, we performed principal component (PC) analysis of CES editing levels. The cell composition was the major factor influencing the observed editing levels, with all the 7 cell types significantly associated with the first 5 PCs. Even if the variance explained by single components is generally low (PC1 ∼ 0.025), our data also revealed 11 biological and pharmacological factors with a significant correlation (p-value < 0.05) with one of the first 5 PCs after correction for cell type composition (Figure 6 and Additional file 1: Table S5).

**Figure 6.**
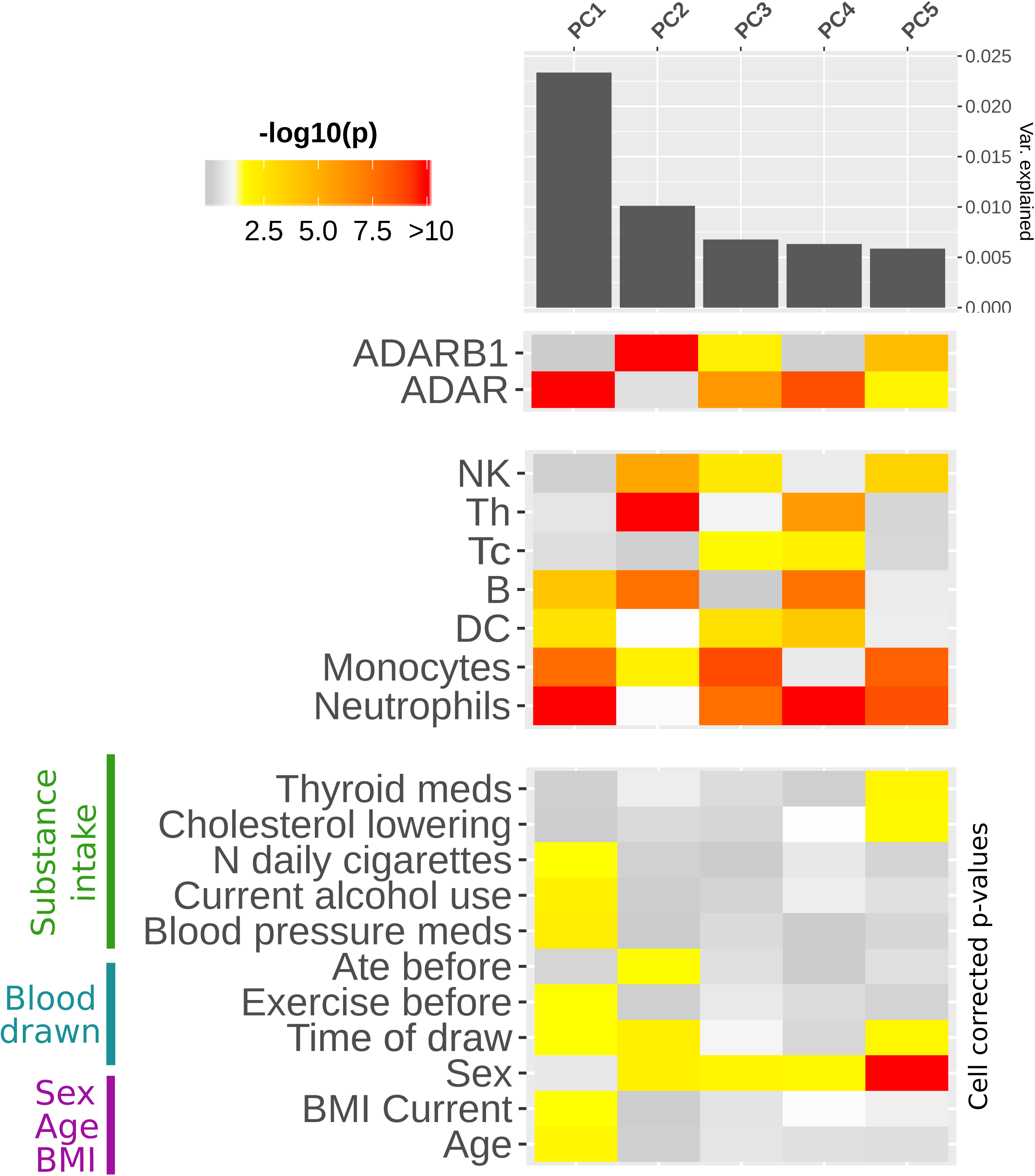
Impact of cell composition, biological and pharmacological factors on PCs of editing levels The heathmap represents strength of association between the first 5 principal components of CESs (PCs) and *ADAR* / *ADARB1* expression (upper panel), 7 cell composition variables (middle panel) and 11 biological / pharmacological variables (lower panel). Only factors showing significant association with at least one of the first 5 PCs are represented. Significant p values (< 0.05) are colored in yellow-red scale, while p value > 0.05 are represented in grey scale. Age, BMI, blood pressure medications, smoke and alcohol all associate with PC1. Also time of blood draw seems to have a small, but consistent effect, on different PCs. For each PC, variance explained is represented by the bar plot in the upper side.

Age and BMI, together with blood pressure medications, smoke and alcohol, seem to be major contributors to editing variability being associated with PC1. Also the time of blood draw seems to have a small, but consistent effect on different PCs. Few variables related to drugs assumption, eating or exercise also emerged with significant association to lower PCs, even if drug intake may be influenced by sex biased distribution (Additional file 1: Table S6). *ADAR* expression level is strongly associated with the first PC, confirming its pivotal role in shaping editing levels variability. Instead, the second PC was associated to expression level for *ADARB1*, but not *ADAR*, suggesting a selective action on a specific group of sites (Figure 6 and Additional file 1: Table S7). Correlation of editing levels for single sites with the first 5 PCs are reported in Additional file 6.

### Identification of genetic variants influencing CES total editing rate

We performed genome wide association analysis between genotyping data of 734,251 SNPs and CES total editing rate to identify SNPs associated with editing levels in human blood (Figure 7a). After variant clumping, our analysis identified a single significant locus on chromosome 7 (rs856554: p-value 3.89e-8), containing the lincRNA gene *LOC730338* (ENSG00000233539) (Figure 7b and Table 3). This locus remains significantly associated with total editing rate after correction for blood cell composition, even if at lower level (Table 3). The SNP rs856554 showed a significant effect on CES total editing rate, while its influence on *ADAR* or *ADARB1* expression was not significant (Figure 7c). Association results for single SNPs with nominal p-value < 0.05 and for loci after variants clumping are reported in Additional file 7. Among genotyped SNPs, there were also 36 known *ADAR* eQTLs, these SNPs explained 5.5% of CES total editing rate variability (p value 3.46e-4). Results of association with CES total editing rate for the known ADAR eQTLs are reported in Additional file 1: Table S8. The effect of the top associated *ADAR* eQTL (rs6699825) on *ADAR* expression and CES total editing rate is represented in Additional file 1: Figure S8.

**Figure 7.**
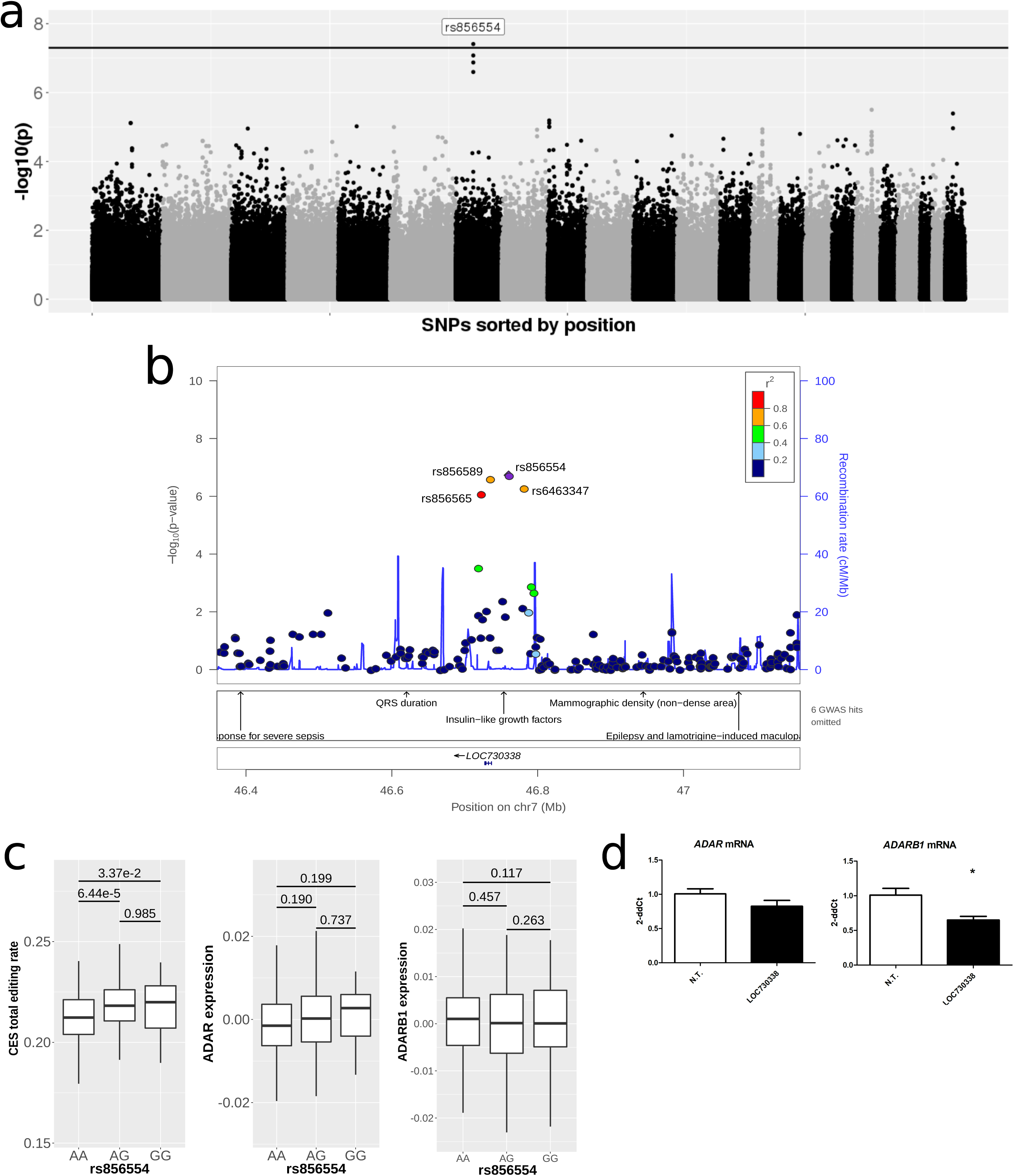
Association study for SNPs and CES total editing rate (a) Manhattan plot representing the association between 573,801 SNPs and CES total editing rate, where black line represents threshold for the top 100 SNPs (p value ∼ 10e-4). (b) Detailed view of genotyped SNPs located in the region at chromosome 7 that showed significant association with CES total editing rate. Known GWAS associations for human phenotypes from GRASP database are reported in the lower panel. (c) The top associated SNP (rs856554) showed a significant effect on global editing level, while no significant correlation was observed with *ADAR* and *ADARB1* expression. (d) Real-time expression analysis of *ADAR* and *ADARB1* mRNA after B-EBV transfection of *LOC730338.* Not transfected cells were used as control samples. Data are reported as 2^-ΔΔ^ct (expression level of control sample is equal to 1) and represent mean values and standard errors obtained from at least 3 independent evaluations. Unpaired t test was used for statistical analysis (*p< 0.05).

**Table 3.**
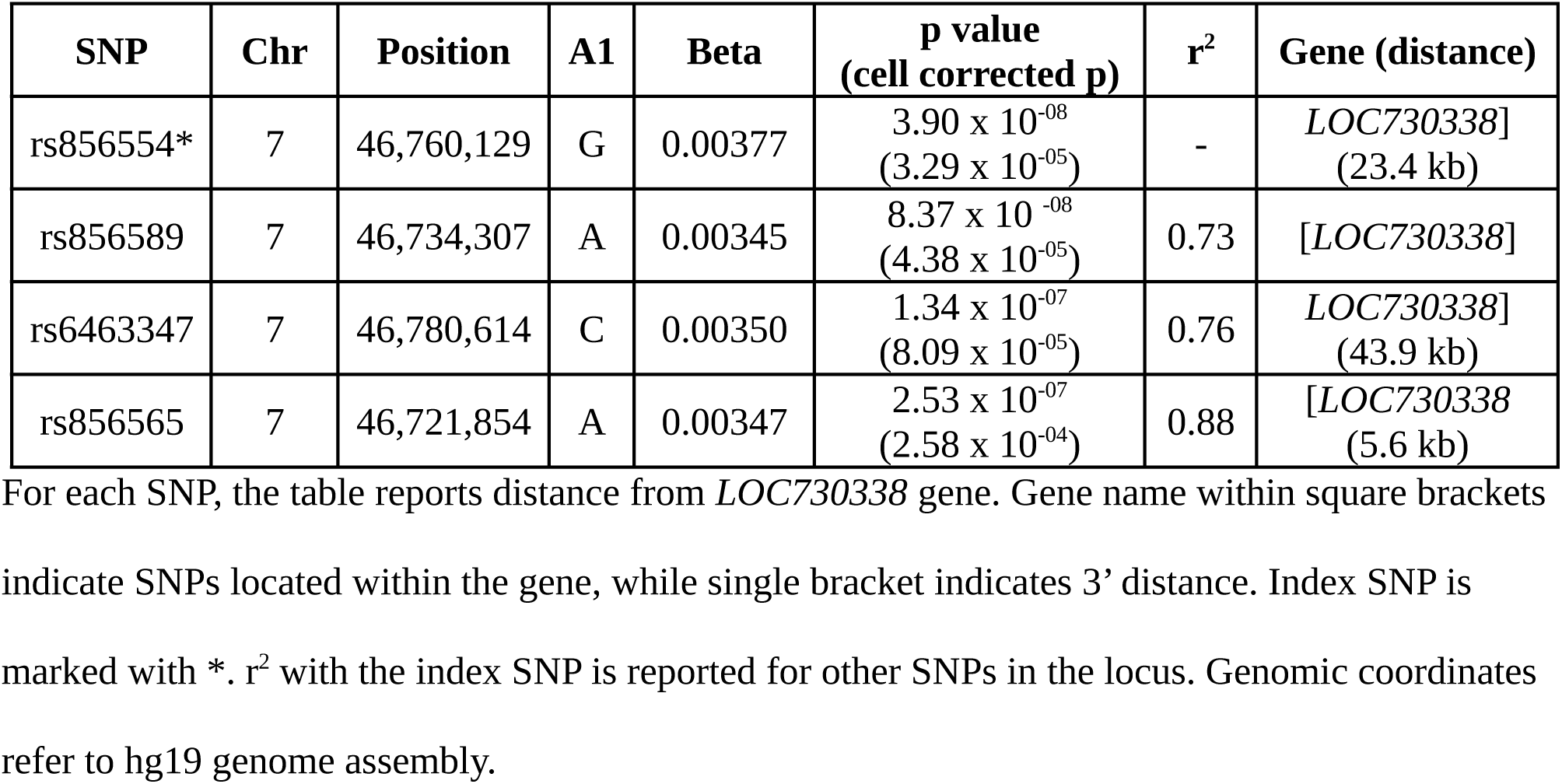
Top 4 SNPs associated to CES total editing rate define a locus on chromosome 7

Since the function of *LOC730338* is unknown, we investigated its possible role in RNA editing, by testing its expression in blood and the effects of its over-expression on ADARs levels in B-EBV cells (Figure 7d). *LOC730338* resulted to be express in blood (data not shown) and its over-expression was associated with a significant down-regulation of *ADARB1* mRNA (Not Transfected cells: 1.00±0.09; *LOC730338* transfected cells: 0.65 ± 0.09; p < 0.05). As concerns ADAR, its expression level was lower in transfected cells (0.85 ± 0.08) than in not transfected (1.00 ± 0.07), but the difference was not statistically significant (p = 0.18).

This result indicates that *LOC730338* might have a limited but noteworthy effect on ADAR enzymes, and it suggests that SNPs inside *LOC730338* could actually affect CES total editing rate by differentially modulating ADAR level.

## Discussion

The process of A-to-I RNA editing has gained increasing attention in recent years, being implicated in multiple aspects of human physiology and, when dysregulated, in human diseases, such as neurological disorders and cancer [2, 9, 24]. Thanks to advances in next-generation sequencing technology, the prevalence and dynamics of “RNA editome” have been recently characterized across many tissues and developmental stages [18, 19, 21, 22]. Overall, more than 2 million editing sites have been described so far, but most of them occur at very low levels in inverted repeat ALU sequences and likely represent random editing with low impact on biological functions [32]. To focus only on those sites that are most likely biologically relevant in human blood, we first selected consistently edited sites (CES) across our dataset of about 450 RNA-Seq samples, resulting in a group of 2,079 sites with at least 5 % editing in at least 100 individuals.

As expected, the majority of these sites is located in inverted repeat ALU sequences [18, 19] that facilitate the formation of a RNA double stranded secondary structure with high affinity for ADAR editing enzymes. Interestingly, nearly 60% of detectable editing sites are located in the 3’UTRs and 37.7% of them fall within a known miRNA binding site. This suggests a potential extensive role of editing process in modulating the miRNA mediated regulation of gene expression in blood [33–35]. Especially, we identified four CES located in conserved miRNA binding sites recognized by conserved miRNAs. Interestingly, three of them regulate the gene *CNPY3*, that might need further investigation.

We identified 22 editing sites located in coding sequences: 12 resulting in synonymous modifications and 10 inducing non-synonymous amino acid changes (re-coding sites). Among the latter, there were well studied re-coding sites, such as the S/G site of *AZIN1* [8], the G/R site of *BLCAP* [36], and the L/R site of *NEIL1* [37]. Their editing levels range from high (75% of *NEIL1*) to medium-low (14% and 16% for *BLCAP* and *AZIN1*, respectively), indicating that both edited and unedited isoforms are needed for the proper function of the tissues. Interestingly, among the re-coding sites, we also detected sites with a high editing level, such as two sites edited at 70% on the small subunit processome component gene (*UTP14C*). It is worthwhile notice that in blood cells 33 editing sites in 3’UTR and intronic regions reach an editing level of more than 90%. Even if we adopted stringent filtering criteria and we observed good concordance in editing levels between our data and REDIportal, we can not completely exclude that extremely high editing levels could results from systematic analysis artifacts. These results prompt for further investigations to understand the actual functional effect of these fully edited sites. Finally, it is of notice that nearby editing sites might correlate in their editing level changes. Correlation is generally strong only for sites closer than 50 nt, but we also detected 5 significant correlations (rho value 0.3 - 0.5) between editing sites on different chromosomes, indicating the possibility of co-regulation mechanisms. Overall, RNA editing process in human blood seems more pervasive than previously reported, prompting for further analyses to understand its biological effects also in healthy subjects.

Further, we investigated the association of genes expression with total editing rate of CES. *ADAR* (encoding ADAR1 enzyme) resulted as the top associated gene and its expression explained about 13% of observed variability, while *ADARB1* (encoding ADAR2 enzyme) was not associated with global editing level even when ALU and non-ALU sites were considered separately. *ADARB2* (encoding ADAR3 protein) is not expressed in blood cells, excluding the possibility that it could have a major negative effect on the editing levels in blood as observed for brain tissues [22]. Thus, ADAR1 emerges as the major contributor to editing process in blood, as already reported for human B cells and other tissues [22, 38], while other ADAR enzymes seem to have only a limited effect. Overall, association analysis revealed 4,719 genes that might have a potential effect on the editing process, strongly enriched for genes involved in the immune system and interferon signaling. This supports the association between genes involved in the inflammatory processes and A-to-I editing in blood cells. Indeed, ADAR1 is present in two main isoforms, a constitutive p110 and an interferon inducible p150 form that is active under an inflammatory response [39]. Moreover, RNA editing, especially ADAR1 activity, is important to modulate innate immunity [40–42]. Modification in the global editing level has been reported after inflammation in mouse and in vitro studies using several inflammatory mediators [43].

When the effect of *ADAR* expression is removed from our analysis, new genes associated with global editing level emerged. These genes are mainly involved in RNA metabolism and ribonucleoprotein complex processing, confirming what has recently been found from a global analysis of GTEx data [22] and strengthening the role of RNA editing complex in RNA processing [38]. Associated genes after *ADAR* correction are strongly enriched for genes encoding for potential ADARs interactors, as revealed by network analysis using data from protein-protein interaction databases. Moreover, associated genes interacting with ADARs mainly encode for RNA binding proteins, as revealed by enrichment analysis, suggesting that they could be involved in RNA recognition or assembly of the editing complex. Network analysis showed that ADAR1 is the main editing enzyme involved in these interactions, confirming its important role in blood samples, compared to the other editing enzymes. We also identified 9 associated genes whose protein products have a direct interaction with ADARs. Among them, proteins encoded by *ELAVL1, RPA1* and *IFI16* emerged as relevant hubs in the network, aggregating most of the interactions directed to ADARs proteins. The stabilizing RNA-binding protein human antigen R (HuR), encoded by *ELAVL1*, has been recently proposed as an ADAR1 interactor involved in the regulation of transcripts stability in human B cells [30, 38]. The observed association between the global editing level and the *ELAVL1* expression strengthens a general role of RNA editing in RNA stability through the modulation of expression of genes involved in RNA metabolism.

Until today, *RPA1* and *IFI16* have never been directly involved in ADARs activity. Our results suggest that they might represent new interesting partners of ADAR1 and that they might help in understand the function and regulation of this key editing enzyme, but larger studies in different cell populations are required to fully understand the impact of these interactions. *RPA1* gene encodes the largest subunit of the heterotrimeric Replication Protein A (RPA) complex, which binds to single-stranded DNA, forming a nucleoprotein complex that is involved in DNA replication, repair, recombination, telomere maintenance and response to DNA damage [44]. ADAR1 presents Z-DNA binding domains, which are not present in the other editing enzymes [45], helping to direct ADAR1 to active transcription sites and to interact with DNA. Thus, the interaction with RPA1 protein might broaden ADAR1 activity also in the field of DNA repair and maintenance. *IFI16*, interferon gamma inducible protein 16, encodes a member of the HIN-200 (hematopoietic interferon-inducible nuclear antigens with 200 amino acid repeats) cytokines family. This protein interacts with p53 and retinoblastoma-1 and localizes to the nucleoplasm and nucleoli [46], where ADAR enzymes are also present. Both IFI16 protein and ADAR1 were associated with response to viral DNA and regulation of immune and interferon signaling responses [46, 47].

RNA editing is known to be a strong tissue-dependent event [22]. Moreover, it has been suggested that the extent of RNA editing may be different among cell types even in the same tissue [23, 48]. In particular, RNA editing events were showed to distribute differently among different cell types in the brain [23]. For this reason, it has been proposed that changes in cellular composition might be responsible for alterations observed in the tissue-wide editing patterns in pathological conditions [49]. In this study, we investigated the relationship between the proportion of different blood cell types (predicted from gene expression data) and the total editing rate of CES. Among the seven different cell types considered, CES total editing rate was positively correlated with the percentage of neutrophils, monocytes, T helper and dendritic cells. These correlations seem mostly mediated by the differential expression of the two ADARs enzymes in the cell populations. The positive correlations observed in neutrophils, monocytes and dendritic cells seem mediated by *ADAR*, whose expression is positive correlated with the percentage of these cell types in blood. Whereas in T helper, editing levels seems mainly mediated by *ADARB1*, whose expression is strongly correlated with the percentage of T helper in blood. This result is also corroborated by the PC analysis, suggesting that the two enzymes have different targets in blood cells. *ADAR* expression is mainly associated with the PC1, supporting its pivotal role in shaping editing levels variability, whereas *ADARB1* with the PC2. Therefore, we could hypothesize a different role of these enzymes in specific cell types (i. e.: T Helper and Natural Killer) and on specific groups of genes. In particular, the specific role of *ADARB1* in T helper deserve further analysis. Overall, this data indicates that cellular composition of the sample should be taken into account carefully to avoid biased results when analyzing editing variations among different groups, such as in case / control studies. However, in our study the amount of the different cells were inferred from expression data [31] and not directly assessed and future replications of these results using direct analysis of purified cells will complete the picture of cell specific editing regulation.

Recently, global editing level has been investigated across tissues and in different species [21, 22] and has also been correlated with the genetic background of human population [30, 50] and with common disease variants [29]. However, the published studies lack a detailed characterization of samples that allows assessing the role of biological and environmental factors.

Relying on on the dataset from [31], containing several demographic, biological and pharmacological variables, we also investigated the potential impact of these external factors on RNA editing process genome-wide. Five variables showed significant correlations with CES total editing rate, namely blood pressure medications, sex, age and body mass index (BMI, current and max). Except for sex, their effect on editing levels seems mainly driven through modulation of *ADAR* expression. Given the strong correlation observed between cell composition, total editing rate and ADARs expression, it is possible that these variables may exert their effect by modulating the proportion of different cell types. However, blood pressure medication, age and current BMI remain correlated also after correction for cellular composition, indicating that they may have a direct effect on the editing levels. Correlation between age and editing was already reported during brain development both in rat [51] and in primates [52] and our data strengthens this correlation also outside the central nervous system. Finally, and for the first time, our study correlated CES editing levels with BMI and blood pressure medications, shedding light on new medical areas in which editing regulation may be involved. A more detailed analysis using principal components of editing levels revealed eleven biological and pharmacological factors significant correlated with PCs even after cell type correction. Age and BMI, together with blood pressure medications, smoke and alcohol, seem to be relevant contributors to editing variability being associated with PC1.

Finally, we analyzed genotyping data to identify SNPs associated with CES total editing rate. Known *ADAR* eQTLs were among the SNPs with the best association p-values and, taken together, they explain about 5% of the observed variation in global editing. This data indicates that the genetic background affects total editing level by modulating ADAR expression level and it must be taken into account when investigating editing regulation in disease studies. A significant association with global editing level in blood was observed for a locus mapping on chromosome 7. This locus contains *LOC730338*, a gene encoding for a long intergenic noncoding RNA (lincRNA). lincRNAs are transcripts longer than 200 nucleotides that have been identified in mammalian genomes mainly by bioinformatics analysis of transcriptomic data. Although thousands of lincRNAs are now validated, the exact function remains unknown for most of them. lincRNAs appear to contribute to the control of gene expression and have a role in cell differentiation and maintenance of cell identity [53]. In *C. elegans,* it has been recently reported that lncRNAs are extensively down-regulated in the absence of ADARs as a result of siRNA generation [54]. The authors suggest that ADARs can interfere with the generation of siRNAs by endogenous RNAi and promote lncRNA expression. *LOC730338* expression cannot be measured in our dataset since it lacks a poly-A tail; therefore, it was not possible to assess if the SNPs associated to total RNA editing rate in the locus are eQTLs of *LOC730338*. However, to go further in understanding the role of *LOC730338* on the editing reaction, we overexpressed its RNA in a lymphoblastic cell line and we showed that it significantly down-regulates *ADARB1* mRNA expression and partially *ADAR* mRNA. Taken together expression results indicate, for the first time, a possible role of *LOC730338* in modulating the expression of ADARs enzymes. Further experimental analysis will be required to identify actual eQTLs for *LOC730338* within the associated genomic locus and understand their possible impact on editing dynamics.

## Conclusion

This study provides a detailed picture of the most consistent RNA editing sites and their variability in human blood. Our results confirm the pivotal role of ADAR1 in the regulation of RNA editing process in blood and suggest new genes, genetic variants, biological and environmental variables that are involved in the RNA editing process. Future studies will be required to confirm and clarify their role and their relationship with the ADAR family enzymes.

### Methods

### Description of data

RNA-Seq raw data (aligned reads) was obtained from NIMH repository, NIMH Study 88 / Site 621, dataset 7 (Levinson RNA Sequencing Data). The original data and samples details are described in [31]. This dataset includes poly(A)+ RNA sequencing and genotyping data from blood samples of 922 subjects, 463 MDD patients and 459 control subjects. The present study focuses only on the 459 controls. Data are provided as aligned reads on hg19 human genome assembly with transcript mapped to RefSeq canonical dataset. Samples are sequenced with a median of 65.6 M reads (31.6 - 258.3), resulting in a median of 14,289 (11,660 - 15,137) detectable genes addressed by at least 10 reads (Additional file 1: Figure S9). Only the 14,961 genes covered with at least 10 reads in at least 100 subjects were considered in the present study for association with editing levels.

A detailed phenotypic description including demographic, pharmacological and biological variables is also included for each subject. Among them, we considered only those relevant in at least 30 subjects and not related to MDD clinical evaluation or socio-economic variables. The 28 variables considered in this study are reported in Additional file 1: Table S2. Moreover, each experiment is annotated with a rich set of technical variables, representing quality metrics of RNA sequencing and characteristics of the blood sample. These also includes 10 cell composition variables, which represent proportion of 10 different cell types as predicted from gene expression data (see supplementary methods in the original paper [31]). In this study, we used the normalized gene expression data provided in [31], determined as residuals of ridge regression of log-transformed read counts with 35 technical and cell composition variables. In this way our analysis of gene expression would not be affected by technical or cell composition biases.

### Assessment of editing levels and selection of consistently edited sites

The original aligned reads were de-duplicated using Picard and the editing levels were then determined genome-wide from BAM files using REDITools v.1.0.4 with the following parameters: −t25 −m20 −c10 −q25 −O5 −l −V0.05 −n0.05. Only sites with a minimum coverage of 10 reads were considered, otherwise their editing level was considered as missing.

To reduce the chance of measuring false-positive editing sites, we selected only sites that met the following criteria: i) sites reported within RefSeq genes by RADAR database [27] and never seen as Single Nucleotide Variants in the human population according to 1000G phase3 and ExAC v.0.3.1; ii) sites occurring in regions were incorrect alignments could have generated artifacts in editing detection were filtered out: known pseudogenes from GENCODE v25; segmental duplication with ≥ 99% identity; single exon genes, that are often retrotransposed genes with high similarity to the corresponding parent gene.

The filtered dataset resulted in 709,184 sites, representing > 75% RADAR editing sites occurring in blood expressed genes. Finally, to provide a picture of most biologically relevant editing events in blood, we decided to focus only on sites with detectable editing levels (at least 5%) in at least 100 subjects (∼20% of total individuals) for subsequent quantitative analysis, resulting in a final dataset of 2,079 sites (consistently edited sites, CES).

### Comparison with REDIportal dataset and overlap with miRNA binding sites

We compared editing levels detected in CES from blood samples with similar data obtained from REDIportal [26]. Editing levels were retrieved directly from REDIportal database, containing RNA editing values calculated from 55 body sites of 150 healthy individuals from GTEx project. Mean editing levels of our 2,079 CES were compared with corresponding data reported for blood tissue in REDI portal. To assess concordance between the two datasets, we calculated concordance correlation coefficient between mean editing values detected in our data and reported in REDIportal blood tissue for the 2,003 overlapping sites.

To assess the overlap between identified CES and miRNA binding sites, we computed the intersection between CES and known miRNA binding sites from TargetScan v.7.2 [55] using bedtools. The analysis was performed separately for broadly conserved, conserved and non-conserved miRNAs and miRNA binding sites, as defined by TargetScan.

### Correlation between editing levels across sites

Using Spearman correlation test, we analyzed correlation of editing levels across the 2,079 CES. Each site was analyzed against all other sites for a total of 4,322,241 tests. FDR correction modified as in [56] was used to account for multiple tests with related variables. Corrplot R package v.0.84 was used to analyze correlation matrices and generate correlation plots.

### Association between CES total editing rate and gene expression

To investigate which genes could influence the editing process, we used robust linear regression (robust v.0.4 R package) to assess the association between gene expression levels and the CES total editing rate in each subject. CES total editing rate for each subject was calculated as in Equation 1.

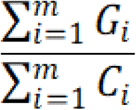

The sum of number of G-containing reads (*Gi*) observed at all CES (*m*), divided by the sum of total reads observed (*Ci*) at all CES.

CES total editing rate was determined also for Alu sites and non-Alu sites, separately. As gene expression levels, we used normalized values provided in [31], where the effect of 35 technical variables, including cell type composition, were regressed out from read counts using ridge regression. In this way, the effect of these variables do not influence subsequent analyses. To choose the set of phenotypic, biological and pharmacological variables to include as covariates in regression analyses, a stepwise model selection by AIC was performed (using stepAIC from MASS R package v.7.3-5). The 6 included variables are indicated in Additional file 1: Table S1. Moreover, since there was a correlation between the variance observed at each editing site and its sequencing coverage for sites with coverage below ∼ 40 X (Additional file 1: Figure S10), the log2 of reads count was also included as covariate in the analysis. The strength of the association was determined by ANOVA test comparing the null (‘background’) model that includes only the set of covariates with the full model (covariates plus normalized expression levels). FDR was used to correct for multiple tests. Subsequently, association analyses were repeated including *ADAR* expression as additional covariate, to remove the effect of *ADAR* expression.

### Gene set enrichment analysis and gene network analysis

The impact on biological functions and cellular pathways of genes found associated with CES total editing rate was investigated using hypergeometric test. We tested the over-representation of pathways among the subset of significant genes at 5% FDR level compared to all expressed genes. Enrichment analysis was performed separately for the following sets from MSigDB v.6.0: cellular pathways from REACTOME and the three main Gene Ontology categories (Cellular Components, GO:CC; Biological Process, GO:BP; Molecular Function, GO:MF). To verify if the proteins encoded by these genes could interact with ADAR proteins, the major enzymes involved in RNA-editing, we explored human protein-protein interaction (PPI) data. First, we created a comprehensive human PPI network combining data from 3 different sources: BioPlex 2.0 [57], BioGRID 3.4.15 [58] and STRING 10.0 [59]. For the BioGRID dataset, only interactions marked as physical were taken in to account, whereas for the STRING dataset only interactions with a combined score above 400 and physical/biochemical evidences were considered. Proteins of the ubiquitin gene family were removed from the network, resulting in a final PPI dataset with 22,913 proteins (nodes) and 833,686 interactions, containing 108 direct interactors of ADARs (ADAR1, ADAR2 and ADAR3 proteins). Among the 376 genes strongly associated with global editing level (FDR < 0.01), we assessed the number of encoded proteins interacting with ADAR1, ADAR2 or one of their first neighbors. To test the significance of these overlap, we performed a random test on the overall set of 14,961 genes addressable in our RNA-Seq data (background genes). We randomly sampled among background genes 1 million groups of N genes (N = 376) and for each simulated group, we counted how many elements interact directly with ADARs or one of their neighbors. Empirical p-value was then calculated as the number of test resulting in an equal or higher number of interactors. Cytoscape v.3.4.0 [60] was used to visualize the PPI network and calculate network related statistics.

### Cell Culture and Co-immunoprecipitation experiments

Epstein-Barr Virus (EBV)-immortalized human B cell lines (B-EBV) were maintained at 37°C, 5% CO2, in RPMI 1640 Medium (Thermo Fisher Scientific), 1% Sodium Pyruvate (Thermo Fisher Scientific), 1% Non-Essential Amino Acid (NEAA, Thermo Fisher Scientific), 15% Fetal Bovine Serum (FBS, Thermo Fisher Scientific), 2mM Glutamine (Thermo Fisher Scientific), 30 U/ml penicillin (Sigma-Aldrich). B-EBV were lysed by sonication in immunoprecipitation buffer (Tris-HCl 50mM pH 7.4, NaCl 300mM, 1% Triton X-100, Protease inhibitors Roche^®^ 1x). The extracts were added to 25µl of Protein G Dynabeads ™ (10007D Invitrogen ^®^ by Thermo Fisher Scientific) coupled with 2µg of mouse anti-ADAR1 (Santa Cruz, cod. sc-73408). After 2h of incubation at 4°C on a rotating wheel, 5 washes with immunoprecipitation buffer were performed. The elution step was carried out with 40µl of Sample buffer 2x and DTT 10x; then the samples were denatured at 95°C for 10 min for the subsequent Western Blot procedure.

During the immunoblot step the following primary antibodies were used 1h at RT: mouse anti-ADAR1 (Santa Cruz, cod. sc-73408) 1:300 in 5% non-fat dry milk in TBST 0,1%; mouse anti-IFI16 (Abcam, cod. Ab55328) 1:500 in 5% non-fat dry milk in TBST 0,1%. The incubation with secondary antibody was performed 1h at RT using the Alkaline Phosphatase (AP)-conjugated anti-mouse secondary antibody 1:10000 in TBST 0,1% (Promega, cod. S372B).

### Identification of cell composition variables correlated with editing levels and ADARs expression

To investigate if cell composition variables could influence editing levels in blood, we considered the 10 cell composition variables described in [31], which represent proportion of 10 different cell types as predicted from gene expression data. After filtering out variables with less than 5% observations (less than 20 subjects), we studied associations between CES total editing rate and 7 cell composition variables (see Additional file 1: Table S2). For these variables, we also analyzed their correlation with *ADAR* and *ADARB1* gene expression levels. Pearson’s product-moment correlation test was used to assess associations.

### Identification of biological factors correlated with editing levels and ADARs expression

To investigate which biological and pharmacological variables could influence editing levels in blood, we studied associations between the 28 biological / pharmacological variables described in Additional file 1: Table S3 and CES total editing rate across subjects, as well as *ADAR* and *ADARB1* expression level. Kruskal-Wallis test, Mann-Whitney-Wilcoxon test and Pearson’s product-moment correlation test were used to assess association for categorical, binary and continuous variables, respectively. We also estimated their effect on CES total editing rate after adjusting for cell composition using the LRT test. This test calculates the impact of a variable of interest on CES total editing rate by comparing a background model containing only covariates, with a full model containing also the variable of interest. In the background model we included only the 4 cell composition variables resulted associated to CES total editing rate (DC, monocytes, neutrophils and Th; see Additional file 1: Table S2), while the full model included also the biological variable of interest. For *ADAR* and *ADARB1* expression, we also estimated the overall effect of the 7 cell composition variables and of biological variables showing an association with the two genes. For each *ADAR* and *ADARB1*, we evaluated the influence of different set of variables using 2 different linear regression model: one including the 7 cell composition variables and the other including also the biological variables. The difference between the two models was assessed using LRT test, to investigate the overall effect of biological variables beyond changes in cell composition.

### Principal component analysis on editing sites

To further investigate the effect of cell composition, biological and pharmacological variables on editing levels in blood, we studied their correlation with the Principal Components of editing levels (PCs). To compute PCs, the missing values of the sites were first imputed using a nonparametric imputation method based on random forest (missForest R package v.1.4 [61]). The PCs were then determined on the complete data using the prcomp R package. To identify the number of PCs to account for, we evaluated the percentage of explained variance by the top 30 PCs, and identified the 5^th^ component as the point at which the explained variance plateaus.

Pearson’s product-moment correlation test was used to assess association for cell composition variables. The association for biological / pharmacological variables was estimated using linear regression model corrected for the 4 cell composition variables resulted associated to CES total editing rate (DC, monocytes, neutrophils and Th; see Additional file 1: Table S2). To identify which editing sites were most correlated with each PC, we analyzed the loadings, that could be interpreted as correlation coefficient between the original variables and components. Moreover, given a high number of sites and low loading values, to deepen the role of each site in the computation of the PCs, we performed the Pearson correlation test. We considered a “moderate” correlation when its absolute value was between 0.3 and 0.5 and the test passed the Bonferroni threshold, while a “weak” correlation was considered when the correlation absolute values ranged between 0 and 0.3 and the respective p-values were significant for Bonferroni correction.

### Association study for SNPs and global editing levels

To identify SNPs associated to global editing level, we analyzed genotyping data and global editing levels in the 459 human blood samples. Starting from genotypes provided in the original dataset [31], we performed quality check removing samples with more than 1% missing genotypes, excessive heterozigosity and PI_HAT > 0.18. Then we removed SNPs with more than 5% missing genotypes, SNPs strongly deviating from Hardy-Weinberg equilibrium (fisher test p-value < 1e-6) and SNPs with a minor allele frequency below 0.01. The final dataset contained 448 individuals and 734,251 SNPs. We used plink v.1.9 linear association analysis with additive model, including as covariates the same 7 variables used for analysis of gene expression (see above, Additional file 1: Table S1) and the first five PCs of genotyping. To identify significant loci associated to global editing level, we performed variant clumping based on the association results, using a 500 kb window and 0.5 R2 threshold. In this way all SNPs in a 500 kb window and with R2 = 0.5 are grouped together around the index SNP, that is the SNP with the lower association p-value. We repeated association analysis including as covariates also the 4 cell composition variables associated to CES total editing rate (DC, monocytes, neutrophils and Th). After association analysis, we used GCTA [62] to evaluate the impact of ADAR known eQTLs on observed global editing levels, using the same set of covariates included for the plink association analysis. This analysis was performed including the 36 known ADAR eQTLs present in our genotyping data.

### Transient transfection of *LOC730338* and ADAR enzymes expression analysis

pTwist-CMV vector containing *LOC30338* sequence between *Not*I and *Bam*HI restriction enzymes, was acquired from Twist Bioscience and used for transient expression of *LOC30338* in B-EBV cells, using lipofectamine 2000 (Thermo Fischer). B-EBV cells were seeded at 100.000 cell/cm2 density in a 6-wells plate. The transfection was performed in Opti-MEM (ThermoFisher Scientific) with a DNA:lipofectamine 1:3 ratio, following manufacturing instructions. Transfected cells were incubated at 37°C for about 24h. After transfection, proteins and RNA were extracted from the cells for further analyses. RNA expression pattern of *ADAR1* and *ADAR2* was analyzed by means of an Applied Biosystems 7500 Real-time PCR system (Applied Biosystems, Foster City, CA, USA). PCR was carried out using TaqMan Universal PCR Master Mix (Applied Biosystems). 25 ng of sample were used in each real-time PCR reaction (TaqMan Gene Expression Assay id probes: *ADAR* (Hs01017596_m1); *ADARB1* (Hs00953724_m1) Applied Biosystems). The expression ratio of target genes in treated sample groups, compared to control group, was calculated using the ΔΔCt method, using *HPRT* (Hs99999909_m1) and *GAPDH* (Hs99999905_m1) geometric means as reference.

## Supporting information

## List of abbreviations

CES: Consistently edited site(s)
GO:MF: Gene Ontology Molecular Function
GO:BP: Gene Ontology Biological Processes
GO:CC: Gene Ontology Cellular Component
eQTL: expression Quantitative Trait Locus

## Declarations

### Ethics approval and consent to participate

Not applicable

### Consent for publication

Not applicable

### Availability of data and material

RNA-Seq raw data and description of subjects are available from NIMH repository, Study 88, but restrictions apply to the availability of these data, which were used under license for the current study, and so are not publicly available.

The datasets supporting the conclusions of this article are included within the article (and its additional files). Editing levels calculated from RNA-Seq are available in the GitHub repository, <u>https://github.com/gedoardo83/RNA_editing_blood</u>.

RADAR database: http://rnaedit.com/

GTEx database: https://www.GTExportal.org/home/

### Authors’ contributions

EG, AB conceived and designed the analysis. EG, CS performed bioinformatics and statistical analysis. MG acquired the data. MG and CM provided intellectual input and conceptual advice and revised the paper. AF, JM performed immuno-precipitation, transfection and expression experiments. EG, AB, CS wrote the paper. All authors read and approved the final manuscript.

## Acknowledgements

Analyzed data come from NIMH Study 88 – Data and biomaterials were provided by Dr. Douglas F. Levinson. This project was supported by National Institutes of Health/National Institute of Mental Health grants 5RC2MH089916 (PI: Douglas F. Levinson, M.D.; Co-investigators: Myrna M. Weissman, Ph.D., James B. Potash, M.D., MPH, Daphne Koller, Ph.D., and Alexander E. Urban, Ph.D.) and 3R01MH090941 (Co-investigator: Daphne Koller, Ph.D.).

## Additional materials

Additional file 1 (pdf). Supplementary tables and figures

Supplementary tables S1-S8. Supplementary Figures S1-S8

Additional file 2 (xls). Detailed statistics for the 2,079 editing sites considered in the study and overlap with miRNA binding sites

Additional file 3 (xls). Complete results of correlation analysis for the 2,079 CES.

Additional file 4 (xls). Complete results of robust regression between CES total editing rate and gene expression levels where the effect of *ADAR* expression was removed.

Additional file 5 (xls). Node properties in the protein-protein interaction network including proteins encoded by genes associated to CES total editing rate (FDR<0.01) and interacting with ADARs or one of their first neighbors.

Additional file 6 (xls). Association of editing sites with principal components of editing.

Additional file 7 (xls). Results of genome-wide association study for CES total editing rate. Association results for single SNPs with nominal p-value < 0.05 and loci identified after variant clumping are reported.

## References

1. Nishikura K. Functions and regulation of RNA editing by ADAR deaminases. Annu Rev Biochem. 2010;79:321–49.

2. Behm M, Öhman M. RNA Editing: A Contributor to Neuronal Dynamics in the Mammalian Brain. Trends Genet. 2016;32:165–75.

3. Bass BL. RNA editing by adenosine deaminases that act on RNA. Annu Rev Biochem. 2002;71:817–46.

4. Orlandi C, Barbon A, Barlati S. Activity regulation of adenosine deaminases acting on RNA (ADARs). Mol Neurobiol. 2012;45:61–75.

5. Hartner JC, Schmittwolf C, Kispert A, Müller AM, Higuchi M, Seeburg PH. Liver disintegration in the mouse embryo caused by deficiency in the RNA-editing enzyme ADAR1. J Biol Chem. 2004;279:4894–902.

6. Higuchi M, Maas S, Single FN, Hartner J, Rozov A, Burnashev N, et al. Point mutation in an AMPA receptor gene rescues lethality in mice deficient in the RNA-editing enzyme ADAR2. Nature. 2000;406:78–81.

7. Wang Q, Khillan J, Gadue P, Nishikura K. Requirement of the RNA editing deaminase ADAR1 gene for embryonic erythropoiesis. Science. 2000;290:1765–8.

8. Chen L, Li Y, Lin CH, Chan THM, Chow RKK, Song Y, et al. Recoding RNA editing of AZIN1 predisposes to hepatocellular carcinoma. Nat Med. 2013;19:209–16.

9. Khermesh K, D'Erchia AM, Barak M, Annese A, Wachtel C, Levanon EY, et al. Reduced levels of protein recoding by A-to-I RNA editing in Alzheimer’s disease. RNA. 2016;22:290–302.

10. Wahlstedt H, Daniel C, Ensterö M, Ohman M. Large-scale mRNA sequencing determines global regulation of RNA editing during brain development. Genome Res. 2009;19:978–86.

11. Sansam CL, Wells KS, Emeson RB. Modulation of RNA editing by functional nucleolar sequestration of ADAR2. Proc Natl Acad Sci U S A. 2003;100:14018–23.

12. Filippini A, Bonini D, Lacoux C, Pacini L, Zingariello M, Sancillo L, et al. Absence of the Fragile X Mental Retardation Protein results in defects of RNA editing of neuronal mRNAs in mouse. RNA Biol. 2017;14:1580–91.

13. Tariq A, Garncarz W, Handl C, Balik A, Pusch O, Jantsch MF. RNA-interacting proteins act as site-specific repressors of ADAR2-mediated RNA editing and fluctuate upon neuronal stimulation. Nucleic Acids Res. 2013;41:2581–93.

14. Garncarz W, Tariq A, Handl C, Pusch O, Jantsch MF. A high-throughput screen to identify enhancers of ADAR-mediated RNA-editing. RNA Biol. 2013;10:192–204.

15. Marcucci R, Brindle J, Paro S, Casadio A, Hempel S, Morrice N, et al. Pin1 and WWP2 regulate GluR2 Q/R site RNA editing by ADAR2 with opposing effects. EMBO J. 2011;30:4211–22.

16. Sommer B, Köhler M, Sprengel R, Seeburg PH. RNA editing in brain controls a determinant of ion flow in glutamate-gated channels. Cell. 1991;67:11–9.

17. Burns CM, Chu H, Rueter SM, Hutchinson LK, Canton H, Sanders-Bush E, et al. Regulation of serotonin-2C receptor G-protein coupling by RNA editing. Nature. 1997;387:303–8.

18. Levanon EY, Eisenberg E, Yelin R, Nemzer S, Hallegger M, Shemesh R, et al. Systematic identification of abundant A-to-I editing sites in the human transcriptome. Nat Biotechnol. 2004;22:1001–5.

19. Li JB, Levanon EY, Yoon J-K, Aach J, Xie B, Leproust E, et al. Genome-wide identification of human RNA editing sites by parallel DNA capturing and sequencing. Science. 2009;324:1210–3.

20. Bazak L, Haviv A, Barak M, Jacob-Hirsch J, Deng P, Zhang R, et al. A-to-I RNA editing occurs at over a hundred million genomic sites, located in a majority of human genes. Genome Res. 2014;24:365–76.

21. Picardi E, Manzari C, Mastropasqua F, Aiello I, Erchia AMD, Pesole G. Profiling RNA editing in human tissues: towards the inosinome Atlas. Nat Publ Gr. 2015;:1–16.

22. Tan MH, Li Q, Shanmugam R, Piskol R, Kohler J, Young AN, et al. Dynamic landscape and regulation of RNA editing in mammals. Nature. 2017;550:249–54.

23. Picardi E, Horner DS, Pesole G. Single-cell transcriptomics reveals specific RNA editing signatures in the human brain. RNA. 2017;23:860–5.

24. Fritzell K, Xu L-D, Lagergren J, Öhman M. ADARs and editing: The role of A-to-I RNA modification in cancer progression. Semin Cell Dev Biol. 2017.

25. Filippini A, Bonini D, La Via L, Barbon A. The Good and the Bad of Glutamate Receptor RNA Editing. Mol Neurobiol. 2016.

26. Picardi E, D'Erchia AM, Lo Giudice C, Pesole G. REDIportal: a comprehensive database of Ato-I RNA editing events in humans. Nucleic Acids Res. 2017;45:D750–7.

27. Ramaswami G, Li JB. RADAR: a rigorously annotated database of A-to-I RNA editing. Nucleic Acids Res. 2014;42 Database issue:D109–13.

28. Gu T, Gatti DM, Srivastava A, Snyder EM, Raghupathy N, Simecek P, et al. Genetic Architectures of Quantitative Variation in RNA Editing Pathways. Genetics. 2016;202:787–98.

29. Franzén O, Ermel R, Sukhavasi K, Jain R, Jain A, Betsholtz C, et al. Global analysis of A-to-I RNA editing reveals association with common disease variants. PeerJ. 2018;6:e4466.

30. Stellos K, Gatsiou A, Stamatelopoulos K, Perisic Matic L, John D, Lunella FF, et al. Adenosineto-inosine RNA editing controls cathepsin S expression in atherosclerosis by enabling HuR-mediated post-transcriptional regulation. Nat Med. 2016;22:1140–50.

31. Mostafavi S, Battle A, Zhu X, Potash JB, Weissman MM, Shi J, et al. Type I interferon signaling genes in recurrent major depression: Increased expression detected by whole-blood RNA sequencing. Mol Psychiatry. 2014;19:1267–74.

32. Ulbricht RJ, Emeson RB. One hundred million adenosine-to-inosine RNA editing sites: hearing through the noise. Bioessays. 2014;36:730–5.

33. Brümmer A, Yang Y, Chan TW, Xiao X. Structure-mediated modulation of mRNA abundance by A-to-I editing. Nat Commun. 2017;8:1255.

34. Borchert GM, Gilmore BL, Spengler RM, Xing Y, Lanier W, Bhattacharya D, et al. Adenosine deamination in human transcripts generates novel microRNA binding sites. Hum Mol Genet. 2009;18:4801–7.

35. Soundararajan R, Stearns TM, Griswold AL, Mehta A, Czachor A, Fukumoto J, et al. Detection of canonical A-to-G editing events at 3' UTRs and microRNA target sites in human lungs using next-generation sequencing. Oncotarget. 2015;6:35726–36.

36. Levanon EY, Hallegger M, Kinar Y, Shemesh R, Djinovic-Carugo K, Rechavi G, et al. Evolutionarily conserved human targets of adenosine to inosine RNA editing. Nucleic Acids Res. 2005;33:1162–8.

37. Yeo J, Goodman RA, Schirle NT, David SS, Beal PA. RNA editing changes the lesion specificity for the DNA repair enzyme NEIL1. Proc Natl Acad Sci U S A. 2010;107:20715–9.

38. Wang IX, So E, Devlin JL, Zhao Y, Wu M, Cheung VG. ADAR regulates RNA editing, transcript stability, and gene expression. Cell Rep. 2013;5:849–60.

39. George CX, John L, Samuel CE. An RNA editor, adenosine deaminase acting on double-stranded RNA (ADAR1). J Interferon Cytokine Res. 2014;34:437–46.

40. Mannion NM, Greenwood SM, Young R, Cox S, Brindle J, Read D, et al. The RNA-editing enzyme ADAR1 controls innate immune responses to RNA. Cell Rep. 2014;9:1482–94.

41. Song C, Sakurai M, Shiromoto Y, Nishikura K. Functions of the RNA Editing Enzyme ADAR1 and Their Relevance to Human Diseases. Genes (Basel). 2016;7:129.

42. Chung H, Calis JJA, Wu X, Sun T, Yu Y, Sarbanes SL, et al. Human ADAR1 Prevents Endogenous RNA from Triggering Translational Shutdown. Cell. 2018;172:811–824.e14.

43. Yang J-H, Luo X, Nie Y, Su Y, Zhao Q, Kabir K, et al. Widespread inosine-containing mRNA in lymphocytes regulated by ADAR1 in response to inflammation. Immunology. 2003;109:15–23.

44. Liu T, Huang J. Replication protein A and more: single-stranded DNA-binding proteins in eukaryotic cells. Acta Biochim Biophys Sin (Shanghai). 2016;48:665–70.

45. Barraud P, Allain FH-T. ADAR proteins: double-stranded RNA and Z-DNA binding domains. Curr Top Microbiol Immunol. 2012;353:35–60.

46. Choubey D, Panchanathan R. IFI16, an amplifier of DNA-damage response: Role in cellular senescence and aging-associated inflammatory diseases. Ageing Res Rev. 2016;28:27–36.

47. Samuel CE. Adenosine deaminases acting on RNA (ADARs) are both antiviral and proviral. Virology. 2011;411:180–93.

48. Leong W-M, Ripen AM, Mirsafian H, Mohamad S Bin, Merican AF. Transcriptogenomics identification and characterization of RNA editing sites in human primary monocytes using high-depth next generation sequencing data. Genomics. 2018.

49. Gal-Mark N, Shallev L, Sweetat S, Barak M, Billy Li J, Levanon EY, et al. Abnormalities in Ato-I RNA editing patterns in CNS injuries correlate with dynamic changes in cell type composition. Sci Rep. 2017;7:43421.

50. Ouyang Z, Ren C, Liu F, An G, Bo X, Shu W. The landscape of the A-to-I RNA editome from 462 human genomes. Sci Rep. 2018;8:12069.

51. Zaidan H, Ramaswami G, Golumbic YN, Sher N, Malik A, Barak M, et al. A-to-I RNA editing in the rat brain is age-dependent, region-specific and sensitive to environmental stress across generations. BMC Genomics. 2018;19:28.

52. Li Z, Bammann H, Li M, Liang H, Yan Z, Phoebe Chen Y-P, et al. Evolutionary and ontogenetic changes in RNA editing in human, chimpanzee, and macaque brains. RNA. 2013;19:1693–702.

53. Cabili MN, Trapnell C, Goff L, Koziol M, Tazon-Vega B, Regev A, et al. Integrative annotation of human large intergenic noncoding RNAs reveals global properties and specific subclasses. Genes Dev. 2011;25:1915–27.

54. Goldstein B, Agranat-Tamir L, Light D, Ben-Naim Zgayer O, Fishman A, Lamm AT. A-to-I RNA editing promotes developmental stage-specific gene and lncRNA expression. Genome Res. 2017;27:462–70.

55. Agarwal V, Bell GW, Nam J-W, Bartel DP. Predicting effective microRNA target sites in mammalian mRNAs. Elife. 2015;4.

56. Benjamini Y, Yekutieli D. The Control of the False Discovery Rate in Multiple Testing under Dependency. The Annals of Statistics. 29:1165–88.

57. Huttlin EL, Bruckner RJ, Paulo JA, Cannon JR, Ting L, Baltier K, et al. Architecture of the human interactome defines protein communities and disease networks. Nature. 2017;545:505–9.

58. Chatr-Aryamontri A, Oughtred R, Boucher L, Rust J, Chang C, Kolas NK, et al. The BioGRID interaction database: 2017 update. Nucleic Acids Res. 2017;45:D369–79.

59. Szklarczyk D, Morris JH, Cook H, Kuhn M, Wyder S, Simonovic M, et al. The STRING database in 2017: quality-controlled protein-protein association networks, made broadly accessible. Nucleic Acids Res. 2017;45:D362–8.

60. Shannon P, Markiel A, Ozier O, Baliga NS, Wang JT, Ramage D, et al. Cytoscape: a software environment for integrated models of biomolecular interaction networks. Genome Res. 2003;13:2498–504.

61. Stekhoven DJ, Bühlmann P. MissForest--non-parametric missing value imputation for mixed-type data. Bioinformatics. 2012;28:112–8.

62. Yang J, Lee SH, Goddard ME, Visscher PM. GCTA: a tool for genome-wide complex trait analysis. Am J Hum Genet. 2011;88:76–82.

