## Supplementary material for "Genome-wide analysis of consistently RNA edited sites in human blood reveals interactions with mRNA processing genes and suggests correlations with cell types and biological variables"

**Table S1.** Summary statistics for the 33 highly edited sites with mean editing level above 0.9

| Site id | Gene | Strand | Region | ALU | Minimum | Mean | Maximum |
|---|---|---|---|---|---|---|---|
| chr18_24179309 | *KCTD1* | - | intronic | no | 0.90 | 1.00 | 1.00 |
| chr6_30453716 | *RANP1* | + | intronic | no | 0.94 | 1.00 | 1.00 |
| chr18_24179333 | *KCTD1* | - | intronic | no | 0.91 | 1.00 | 1.00 |
| chr18_24179324 | *KCTD1* | - | intronic | no | 0.91 | 1.00 | 1.00 |
| chr9_90140405 | *DAPK1* | + | intronic | no | 0.92 | 1.00 | 1.00 |
| chr6_132140917 | *ENPP1* | + | intronic | no | 0.90 | 1.00 | 1.00 |
| chr15_25154978 | *SNRPN* | + | intronic | no | 0.30 | 1.00 | 1.00 |
| chr16_27278365 | *NSMCE1* | - | intronic | no | 0.90 | 1.00 | 1.00 |
| chr3_44619818 | *RP11-944L7.4* | - | intronic | no | 0.90 | 1.00 | 1.00 |
| chr3_183868236 | *EIF2B5* | + | intronic | no | 0.93 | 0.99 | 1.00 |
| chr20_3276419 | *C20orf194* | - | ncRNA | no | 0.92 | 0.99 | 1.00 |
| chr5_130994333 | *FNIP1* | - | intronic | no | 0.82 | 0.99 | 1.00 |
| chr7_38351411 | *TARP* | - | intronic | no | 0.90 | 0.99 | 1.00 |
| chr3_44620372 | *RP11-944L7.4* | - | intronic | no | 0.82 | 0.99 | 1.00 |
| chr22_42777186 | *NFAM1* | - | 3UTR | no | 0.87 | 0.98 | 1.00 |
| chr5_130994309 | *FNIP1* | - | intronic | no | 0.80 | 0.98 | 1.00 |
| chr3_44620354 | *RP11-944L7.4* | - | intronic | no | 0.82 | 0.98 | 1.00 |
| chr7_38351422 | *TARP* | - | intronic | no | 0.80 | 0.98 | 1.00 |
| chr10_101992308 | *CWF19L1* | - | 3UTR | yes | 0.80 | 0.98 | 1.00 |
| chrX_70640516 | *BCYRN1* | - | intronic | no | 0.75 | 0.97 | 1.00 |
| chr12_98943033 | *TMPO* | + | 3UTR | yes | 0.76 | 0.96 | 1.00 |
| chr5_130994344 | *FNIP1* | - | intronic | no | 0.77 | 0.96 | 1.00 |
| chr19_56150729 | *ZNF581* | + | intronic | yes | 0.69 | 0.96 | 1.00 |
| chr19_8557875 | *PRAM1* | - | intronic | yes | 0.80 | 0.95 | 1.00 |
| chr1_160966434 | *F11R* | - | 3UTR | yes | 0.79 | 0.95 | 1.00 |
| chr6_31640793 | *CSNK2B* | + | 3UTR | yes | 0.69 | 0.95 | 1.00 |
| chr15_45025366 | *TRIM69* | + | intronic | yes | 0.7 | 0.95 | 1.00 |
| chr7_128293538 | *LINC01000* | + | ncRNA | no | 0.77 | 0.94 | 1.00 |
| chr14_20834467 | *TEP1* | - | 3UTR | no | 0.67 | 0.93 | 1.00 |

| | | | | | | | |
|---|---|---|---|---|---|---|---|
| chr22_42777158 | *NFAM1* | - | 3UTR | no | 0.69 | 0.92 | 1.00 |
| chr22_39350614 | *APOBEC3A* | + | intronic | no | 0.68 | 0.92 | 1.00 |
| chr19_15772035 | *CYP4F3* | + | 3UTR | yes | 0.66 | 0.91 | 1.00 |
| chr15_90375494 | *C15orf38*<br>*AP3S2* | - | 3UTR | yes | 0.70 | 0.90 | 1.00 |

**Table S2.** Cell composition variables associated to CES total editing rate and *ADAR* / *ADARB1* expression level.
The table reports number of subjects with non-zero value and results of association between cell composition variables and CES total editing rate or *ADAR* / *ADARB1* expression level (TPM). Pearson's product-moment correlation test were used to assess association. Pearson r coefficient is also reported. Variables measurable in less than 20 individuals (5% of total sample) were not analysed. Th = T helper, Tc = T cytotoxic, Tc_act = activated T cells, B = B linfocytes, NK = natural killer linfocytes, NK_act = activated natural killer linfocytes, mono = monocytes, DC = dendritic cells, DC_act = activated dendritic cells, neutro = neutrophils.

| Cell variable | Tested variable | N. of subject with non zero value | P val | r |
|---|---|---|---|---|
| neutro | Total editing rate | 452 | 1.78E-16 | 0.374 |
| neutro | ADAR | 452 | 7.31E-11 | 0.300 |
| neutro | ADARB1 | 452 | 7.24E-19 | -0.401 |
| mono | Total editing rate | 450 | 9.52E-07 | 0.229 |
| mono | ADAR | 450 | 5.38E-02 | 0.091 |
| mono | ADARB1 | 450 | 7.30E-05 | -0.186 |
| Th | Total editing rate | 452 | 3.46E-05 | 0.193 |
| Th | ADAR | 452 | 2.14E-02 | -0.108 |
| Th | ADARB1 | 452 | 3.69E-18 | 0.393 |
| DC | Total editing rate | 104 | 1.56E-04 | 0.362 |
| DC | ADAR | 104 | 6.16E-02 | 0.184 |
| DC | ADARB1 | 104 | 3.08E-01 | -0.101 |
| NK | Total editing rate | 407 | 2.87E-01 | 0.053 |
| NK | ADAR | 407 | 6.03E-03 | 0.136 |
| NK | ADARB1 | 407 | 7.96E-02 | 0.087 |
| Tc | Total editing rate | 50 | 2.91E-01 | 0.152 |
| Tc | ADAR | 50 | 6.91E-01 | -0.058 |
| Tc | ADARB1 | 50 | 6.54E-01 | -0.065 |
| B | Total editing rate | 389 | 7.79E-01 | 0.0143 |
| B | ADAR | 389 | 3.74E-02 | -0.106 |
| B | ADARB1 | 389 | 3.80E-21 | 0.454 |
| DC_act | Total editing rate | 11 | | |
| DC_act | ADAR | 11 | | |
| DC_act | ADARB1 | 11 | | |
| NK_act | Total editing rate | 8 | | |
| NK_act | ADAR | 8 | | |
| NK_act | ADARB1 | 8 | | |
| Tc_act | Total editing rate | 1 | | |
| Tc_act | ADAR | 1 | | |
| Tc_act | ADARB1 | 1 | | |

**Table S3.** Phenotypic / pharmacological variables considered in the study.
The table reports the 28 variables included in the study, extracted from the original dataset described in [29].

| Variable | Explanation | Included as covariate |
|---|---|---|
| Age | Age at recruitment | |
| BMI current | BMI at the time of recruitment (self-report) | x |
| BMI max | Maximum BMI lifetime (self-report) | |
| Region | Region of country | |
| Sex | Sex | x |
| Ate before | Ate before blood draw (self-report) | |
| Exercise before | Exercised before blood draw (self-report) | |
| Smoke before | Smoked before blood draw (self-report) | |
| Time of draw | Time blood was drawn converted to numeric eg, 2:30 pm = 14.5 | |
| All drugs | Used any drug (cannabis, cocaine, hallucinogens, stimulants) | |
| Cannabis | Used cannabis | |
| Cocaine | Used cocaine | |
| Hallucinogens | Used hallucinogens | |
| Stimulants | Used stimulants | |
| ACE inhibitor | Currently use ACE inhibitors | x |
| Blood pressure meds | Currently use blood pressure medications | x |
| Oral birth-control pill | Currently use oral birth control pill | x |
| Cholesterol lowering | Currently use cholesterol lowering medications | |
| Anti histamine | Currently use anti histamine medications | |
| Thyroid meds | Currently use thyroid hormone supplementation | x |
| Proton-pump inhibitor | Currently use proton pump inhibitors | |
| None treatments | None medical treatments | |
| Alcohol abuse | Ever had alcohol abuse | |
| Current alcohol use | Level of current alcohol use | |
| 100 cigs lifetime | Ever smoked 100 cigarettes lifetime | |
| Currently smoke | Currently smoke | |
| N daily cigarettes | Current typical number of cigarettes smoked daily | |
| Fager life total | Total Fagerstrom score, worst lifetime period | |

**Table S4.** Biological / pharmacological factors associated to CES total editing rate and ADARs expression

The table reports association results between biological / pharmacological factors and CES total editing rate or *ADAR* / *ADARB1* expression level (calculated as Transcript per Million). Kruskal-Wallis test, Mann-Whitney-Wilcoxon test and Pearson's product-moment correlation test were used to assess association for categorical, binary and continuous variables, respectively. For continuous variables Pearson r coefficient is also reported. We also calculated a cell adjusted P value, that represents significance of LRT test for impact of each variable after accounting for cell factors associated to CES total editing rate (neutrophils, monocytes, DC, Th). Variables are sorted based on P value of association with total editing rate.

| Variable | Tested association | P value | Cell adjusted P value | r |
|---|---|---|---|---|
| Blood pressure meds | Total editing rate | 3.95E-04 | 8.93E-04 | |
| Blood pressure meds | ADAR | 5.02E-04 | 1.97E-02 | |
| Blood pressure meds | ADARB1 | 9.62E-01 | 4.52E-01 | |
| BMI current | Total editing rate | 1.68E-03 | 4.77E-02 | -0.148 |
| BMI current | ADAR | 3.68E-03 | 2.29E-02 | -0.137 |
| BMI current | ADARB1 | 2.95E-01 | 9.31E-02 | -0.050 |
| Age | Total editing rate | 3.62E-03 | 7.57E-03 | -0.137 |
| Age | ADAR | 1.47E-03 | 5.29E-03 | -0.149 |
| Age | ADARB1 | 2.99E-01 | 1.17E-01 | -0.049 |
| BMI max | Total editing rate | 4.64E-03 | 7.44E-02 | -0.134 |
| BMI max | ADAR | 6.49E-03 | 3.18E-02 | -0.128 |
| BMI max | ADARB1 | 3.32E-01 | 1.24E-01 | -0.046 |
| Sex | Total editing rate | 4.12E-02 | 3.53E-01 | |
| Sex | ADAR | 2.23E-01 | 3.93E-02 | |
| Sex | ADARB1 | 1.72E-01 | 2.52E-02 | |
| Oral birth-control pill | Total editing rate | 5.98E-02 | 1.54E-01 | |
| Oral birth-control pill | ADAR | 1.75E-01 | 5.85E-01 | |
| Oral birth-control pill | ADARB1 | 5.95E-01 | 3.47E-01 | |
| Alcohol abuse | Total editing rate | 6.96E-02 | 4.21E-01 | |
| Alcohol abuse | ADAR | 1.58E-01 | 2.97E-01 | |
| Alcohol abuse | ADARB1 | 9.82E-01 | 4.42E-01 | |
| Cocaine | Total editing rate | 1.54E-01 | 9.01E-01 | |
| Cocaine | ADAR | 7.78E-01 | 3.81E-01 | |
| Cocaine | ADARB1 | 5.46E-01 | 6.87E-01 | |
| ACE inhibitor | Total editing rate | 2.37E-01 | 6.04E-01 | |
| ACE inhibitor | ADAR | 3.93E-01 | 8.91E-01 | |
| ACE inhibitor | ADARB1 | 4.29E-01 | 9.57E-01 | |
| Region | Total editing rate | 2.46E-01 | 1.09E-01 | |
| Region | ADAR | 7.79E-01 | 7.07E-01 | |

| | | | | |
|---|---|---|---|---|
| Region | ADARB1 | 6.10E-01 | 7.24E-01 | |
| Hallucinogens | Total editing rate | 2.46E-01 | 7.30E-01 | |
| Hallucinogens | ADAR | 2.51E-01 | 5.18E-02 | |
| Hallucinogens | ADARB1 | 5.94E-02 | 2.29E-01 | |
| Cholesterol lowering | Total editing rate | 2.51E-01 | 5.59E-01 | |
| Cholesterol lowering | ADAR | 6.33E-01 | 8.00E-01 | |
| Cholesterol lowering | ADARB1 | 8.66E-01 | 9.84E-01 | |
| None treatments | Total editing rate | 3.20E-01 | 4.42E-01 | |
| None treatments | ADAR | 8.89E-01 | 7.14E-01 | |
| None treatments | ADARB1 | 2.39E-01 | 2.87E-01 | |
| 100 cigs lifetime | Total editing rate | 3.63E-01 | 7.39E-01 | |
| 100 cigs lifetime | ADAR | 3.68E-01 | 9.71E-01 | |
| 100 cigs lifetime | ADARB1 | 9.65E-01 | 9.12E-02 | |
| All drugs | Total editing rate | 3.91E-01 | 4.61E-01 | |
| All drugs | ADAR | 4.69E-01 | 2.65E-01 | |
| All drugs | ADARB1 | 3.49E-01 | 3.49E-01 | |
| Cannabis | Total editing rate | 4.53E-01 | 5.51E-01 | |
| Cannabis | ADAR | 6.76E-01 | 4.31E-01 | |
| Cannabis | ADARB1 | 1.64E-01 | 1.81E-01 | |
| Current alcohol use | Total editing rate | 8.32E-02 | 8.03E-02 | |
| Current alcohol use | ADAR | 4.44E-01 | 1.17E-01 | |
| Current alcohol use | ADARB1 | 3.62E-01 | 5.83E-01 | |
| Stimulants | Total editing rate | 5.42E-01 | 9.65E-01 | |
| Stimulants | ADAR | 8.95E-01 | 3.71E-01 | |
| Stimulants | ADARB1 | 1.64E-01 | 4.86E-01 | |
| Time of draw | Total editing rate | 6.02E-01 | 4.66E-01 | -0.025 |
| Time of draw | ADAR | 6.12E-01 | 8.23E-01 | -0.024 |
| Time of draw | ADARB1 | 9.37E-03 | 4.82E-02 | 0.122 |
| Thyroid meds | Total editing rate | 6.08E-01 | 7.65E-01 | |
| Thyroid meds | ADAR | 4.98E-01 | 3.49E-01 | |
| Thyroid meds | ADARB1 | 1.12E-02 | 4.17E-02 | |
| N daily cigarettes | Total editing rate | 6.17E-01 | 4.12E-01 | 0.024 |
| N daily cigarettes | ADAR | 9.46E-01 | 2.70E-01 | -0.003 |
| N daily cigarettes | ADARB1 | 1.49E-01 | 9.40E-01 | 0.068 |
| Fager life total | Total editing rate | 6.26E-01 | 4.91E-01 | -0.023 |
| Fager life total | ADAR | 7.35E-02 | 2.49E-01 | -0.084 |
| Fager life total | ADARB1 | 5.62E-01 | 5.70E-02 | -0.027 |
| Ate before | Total editing rate | 6.71E-01 | 5.66E-01 | |
| Ate before | ADAR | 3.23E-01 | 3.60E-01 | |

| | | | |
|---|---|---|---|
| Ate before | ADARB1 | 4.92E-02 | 1.42E-02 |
| Smoke before | Total editing rate | 7.02E-01 | 3.59E-01 |
| Smoke before | ADAR | 8.21E-01 | 5.42E-01 |
| Smoke before | ADARB1 | 8.97E-02 | 6.96E-01 |
| Currently smoke | Total editing rate | 7.34E-01 | 4.13E-01 |
| Currently smoke | ADAR | 8.97E-01 | 5.84E-02 |
| Currently smoke | ADARB1 | 9.26E-02 | 8.89E-01 |
| Anti histamine | Total editing rate | 7.44E-01 | 2.75E-01 |
| Anti histamine | ADAR | 6.67E-01 | 7.67E-01 |
| Anti histamine | ADARB1 | 8.09E-01 | 9.46E-01 |
| Exercise before | Total editing rate | 9.50E-01 | 2.53E-01 |
| Exercise before | ADAR | 1.00E+00 | 2.59E-01 |
| Exercise before | ADARB1 | 8.94E-01 | 8.78E-01 |
| Proton-pump inhibitor | Total editing rate | 9.60E-01 | 1.08E-01 |
| Proton-pump inhibitor | ADAR | 6.29E-01 | 1.44E-01 |
| Proton-pump inhibitor | ADARB1 | 1.77E-01 | 3.44E-02 |

**Table S5.** Association of cell composition variables, biological and pharmacological variables with the top 5 editing principal components.
For cell types variables, the table reports p-value for association with the first 5 PCs, calculated using Pearson's product-moment correlation test. For biological / pharmacological variables we reported the p-value for association after correcting for the 4 cell variables associated to CES total editing rate (neutrophils, monocytes, DC, Th; see Table S2). Variable are sorted based on association with PC1.

| **Cell composition variables** | | | | | |
|---|---|---|---|---|---|
| **variable** | **PC1** | **PC2** | **PC3** | **PC4** | **PC5** |
| neutrophils | 7.82E-44 | 7.50E-02 | 2.02E-08 | 3.06E-16 | 1.93E-09 |
| monocytes | 1.71E-08 | 9.55E-03 | 1.45E-09 | 1.92E-01 | 6.24E-09 |
| B | 9.70E-05 | 2.82E-08 | 9.61E-01 | 3.04E-08 | 1.77E-01 |
| DC | 1.91E-03 | 6.59E-02 | 1.87E-03 | 1.37E-04 | 1.62E-01 |
| Tc | 3.60E-01 | 7.66E-01 | 2.37E-02 | 9.86E-03 | 5.33E-01 |
| Th | 2.51E-01 | 3.31E-20 | 1.14E-01 | 1.13E-06 | 5.45E-01 |
| NK | 7.83E-01 | 3.55E-06 | 3.89E-03 | 1.81E-01 | 4.04E-04 |
| **Biological / pharmacological variables after correction for cell composition** | | | | | |
| **variable** | **PC1** | **PC2** | **PC3** | **PC4** | **PC5** |
| Blood pressure meds | 7.45E-03 | 9.56E-01 | 4.27E-01 | 9.58E-01 | 5.22E-01 |
| Current alcohol use | 9.31E-03 | 9.40E-01 | 6.60E-01 | 1.40E-01 | 2.92E-01 |
| Age | 1.70E-02 | 7.65E-01 | 2.15E-01 | 2.91E-01 | 3.38E-01 |
| Time of draw | 4.24E-02 | 9.41E-03 | 8.84E-02 | 5.11E-01 | 1.94E-02 |
| Exercise before | 4.29E-02 | 8.35E-01 | 1.75E-01 | 3.69E-01 | 6.43E-01 |
| BMI current | 4.73E-02 | 8.91E-01 | 2.35E-01 | 5.84E-02 | 1.21E-01 |
| N daily cigarettes | 4.77E-02 | 7.17E-01 | 9.74E-01 | 1.88E-01 | 5.87E-01 |
| BMI max | 7.26E-02 | 8.71E-01 | 3.88E-01 | 7.99E-02 | 1.63E-01 |
| Currently smoke | 7.65E-02 | 8.45E-01 | 6.60E-01 | 1.82E-01 | 2.30E-01 |
| Sex | 1.90E-01 | 1.01E-02 | 1.72E-02 | 2.13E-02 | 5.39E-11 |
| Smoke before | 1.94E-01 | 4.18E-01 | 7.86E-01 | 5.89E-01 | 3.21E-01 |
| Alcohol abuse | 2.31E-01 | 3.57E-01 | 1.94E-01 | 3.80E-01 | 6.49E-01 |
| Proton-pump inhibitor | 2.40E-01 | 5.29E-01 | 2.94E-01 | 9.63E-01 | 7.58E-01 |
| All drugs | 2.82E-01 | 9.84E-01 | 3.55E-01 | 7.33E-02 | 4.26E-01 |
| Region | 3.07E-01 | 7.86E-01 | 9.49E-01 | 1.34E-01 | 6.46E-01 |
| Oral birth-control pill | 3.90E-01 | 4.49E-01 | 3.80E-01 | 1.75E-01 | 7.55E-01 |
| Cannabis | 4.53E-01 | 8.98E-01 | 4.39E-01 | 1.97E-01 | 4.84E-01 |
| 100 cigs lifetime | 4.55E-01 | 2.12E-01 | 7.28E-01 | 3.55E-01 | 4.33E-01 |
| Ate before | 5.85E-01 | 3.52E-02 | 3.19E-01 | 9.69E-01 | 3.16E-01 |
| Anti histamine | 6.24E-01 | 4.96E-01 | 8.12E-01 | 8.80E-01 | 9.66E-01 |
| ACE inhibitor | 6.50E-01 | 3.93E-01 | 5.79E-01 | 5.64E-01 | 1.91E-01 |
| Stimulants | 6.97E-01 | 7.34E-01 | 5.63E-01 | 8.30E-01 | 2.69E-01 |

| | | | | | |
|---|---|---|---|---|---|
| Thyroid meds | 7.64E-01 | 1.42E-01 | 3.82E-01 | 8.34E-01 | 1.80E-02 |
| None treatments | 8.39E-01 | 4.25E-01 | 8.03E-01 | 8.55E-01 | 6.62E-01 |
| Cholesterol lowering | 9.11E-01 | 4.22E-01 | 5.33E-01 | 5.42E-02 | 2.42E-02 |
| Cocaine | 9.15E-01 | 6.00E-01 | 4.84E-01 | 4.46E-01 | 4.79E-01 |
| Hallucinogens | 9.94E-01 | 1.78E-01 | 1.43E-01 | 3.85E-01 | 4.62E-01 |

**Table S6.** Association of drug an medication intake variables with sex of subjects. Association was analyzed by Chi-Square test.

| Variable | P-value |
|---|---|
| Oral birth-control pill meds | 3.00E-06 |
| Alcohol abuse | 4.00E-05 |
| Cholesterol lowering | 0.001 |
| Thyroid meds | 0.005 |
| Current alcohol use | 0.02 |
| All drugs | 0.2 |
| Hallucinogens | 0.2 |
| None treatments | 0.2 |
| 100 cigs lifetime | 0.2 |
| Blood pressure meds | 0.3 |
| N daily cigarettes | 0.3 |
| Stimulants | 0.3 |
| Cocaine | 0.4 |
| ACE Inhibitor | 0.5 |
| Cannabis | 0.5 |
| Smoke before | 0.6 |
| Proton-pump inhibitor | 0.7 |
| Currently smoke | 0.8 |
| Anti histamine | 0.9 |

**Table S7.** Association of ADAR and ADARB1 expression level with the top 5 editing principal components.
The table reports p-value for association of ADAR and ADARB1 expression level with the first 5 PCs, calculated using Pearson's product-moment correlation test.

| Gene | PC1 | PC2 | PC3 | PC4 | PC5 |
|---|---|---|---|---|---|
| ADAR | 3.07E-58 | 4.66E-02 | 4.09E-07 | 1.61E-09 | 3.17E-03 |
| ADARB1 | 1.41E-01 | 2.10E-42 | 1.86E-03 | 1.13E-01 | 1.32E-05 |

**Table S8.** Results of association with CES total editing rate for the known ADAR eQTLs. The rank among the 734,251 tested SNPs is reported.

| Rank | SNP | A1 | BETA | P-value |
|---|---|---|---|---|
| 58 | rs903323 | A | -2.47E-03 | 4.19E-05 |
| 59 | rs6699825 | G | -2.47E-03 | 4.19E-05 |
| 64 | rs9426830 | G | -2.45E-03 | 4.76E-05 |
| 124 | rs1127313 | A | -2.30E-03 | 1.22E-04 |
| 154 | rs1127311 | A | -2.28E-03 | 1.44E-04 |
| 480 | rs9427108 | A | -2.09E-03 | 5.58E-04 |
| 658 | rs2335230 | C | 2.61E-03 | 8.04E-04 |
| 1032 | rs9427114 | G | -1.93E-03 | 1.33E-03 |
| 3052 | rs9427097 | C | -2.24E-03 | 4.03E-03 |
| 4316 | rs11264248 | G | 1.66E-03 | 5.74E-03 |
| 4319 | rs7547072 | A | -1.66E-03 | 5.75E-03 |
| 15375 | rs1876304 | A | 1.68E-03 | 2.09E-02 |
| 63347 | rs2072660 | A | 1.23E-03 | 8.67E-02 |
| 87416 | rs7556080 | A | 1.07E-03 | 1.20E-01 |
| 136823 | rs2131902 | G | 8.91E-04 | 1.87E-01 |
| 168738 | rs1127309 | A | 8.18E-04 | 2.30E-01 |
| 170542 | rs9426823 | A | 8.09E-04 | 2.33E-01 |
| 172939 | rs2229857 | A | 8.05E-04 | 2.36E-01 |
| 174891 | rs6656743 | T | 7.98E-04 | 2.39E-01 |
| 174972 | rs7534678 | A | 7.96E-04 | 2.39E-01 |
| 210425 | rs11264235 | A | 7.22E-04 | 2.87E-01 |
| 211471 | rs7554577 | A | 6.71E-04 | 2.89E-01 |
| 219063 | rs1127314 | G | 7.02E-04 | 2.99E-01 |
| 241948 | rs11264222 | A | 6.64E-04 | 3.30E-01 |
| 295150 | rs884617 | A | 5.47E-04 | 4.03E-01 |
| 319610 | rs6696760 | A | 6.89E-04 | 4.36E-01 |
| 326377 | rs2172706 | G | -5.56E-04 | 4.45E-01 |
| 414576 | rs4845617 | A | -3.63E-04 | 5.65E-01 |
| 420648 | rs10908431 | G | 3.71E-04 | 5.73E-01 |
| 506659 | rs2297607 | G | -2.95E-04 | 6.90E-01 |
| 530232 | rs1194587 | G | 2.14E-04 | 7.22E-01 |
| 531916 | rs952146 | G | -2.27E-04 | 7.25E-01 |
| 534952 | rs3738032 | A | -2.57E-04 | 7.29E-01 |
| 575493 | rs2988721 | A | -1.92E-04 | 7.84E-01 |
| 581493 | rs10908835 | G | -1.70E-04 | 7.92E-01 |
| 668056 | rs2274988 | A | -7.97E-05 | 9.10E-01 |

**Figure S1.** Measurable editing sites in our dataset cover most sites in blood expressed genes
We calculated the fraction of total editing sites reported from RADAR that have adequate coverage (at least 10X) in our dataset. Based on expression data from GTeX v7 we calculated this fraction stratified based on most expressed genes in whole blood. Our dataset covers > 75% of total RADAR sites in the top 5,000 genes expressed in whole blood, supporting our ability to investigate editing events in this tissue.

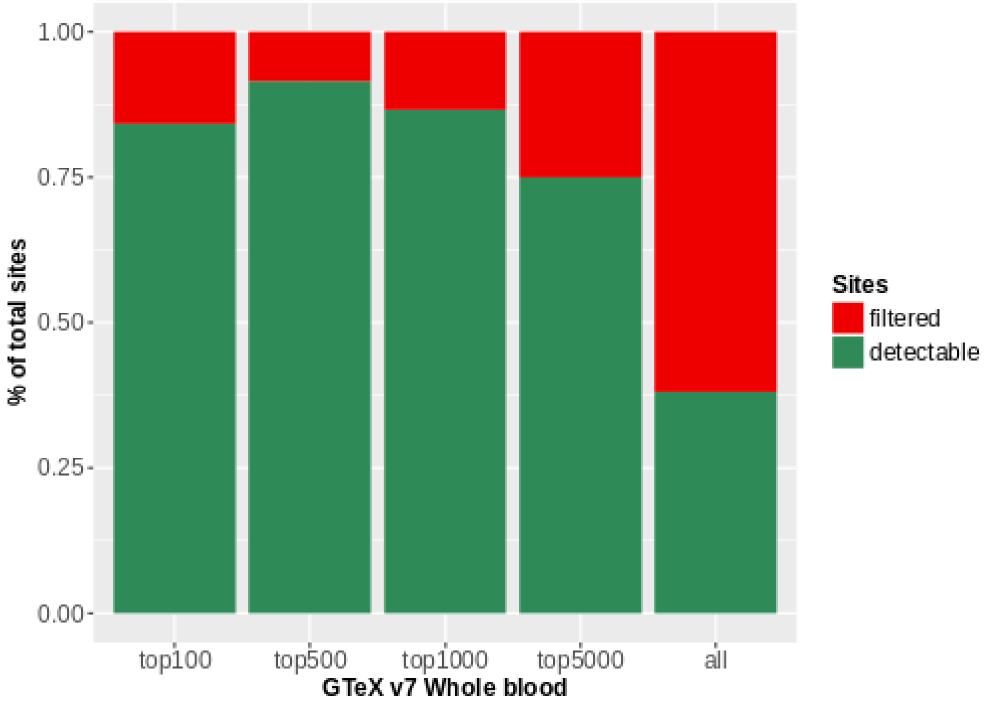

**Figure S2.** Distribution of sites with detectable editing

Considering the 709,184 sites addressable in our dataset, we calculated for each one the number of individuals with detectable editing level. The distribution of number of sites detectable in at least N samples is represented in the plot. We found that 691,304 sites have no detectable editing and most sites have a detectable editing level only in a small fraction of samples. The red dot indicates the number of sites with detectable editing level in at least 100 individuals.

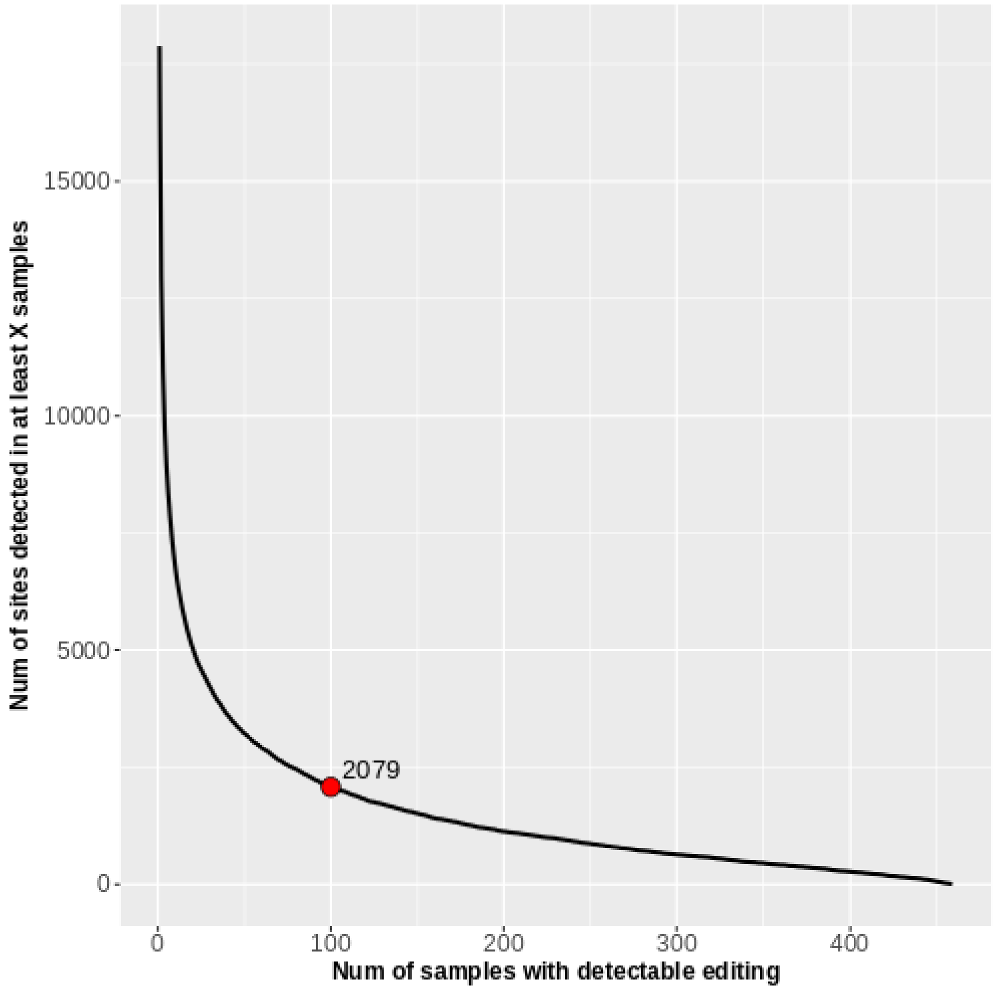

**Figure S3.** Concordance between editing levels observed in our study and REDIPortal
We calculated concordance correlation coefficient (CCC) between mean editing values for the 2,003 overlapping sites detected in our data (Mean) and also reported by REDIPortal (Mean REDI). The blue dashed line represents the fitted linear model, while red line represents linear model for perfect concordance.

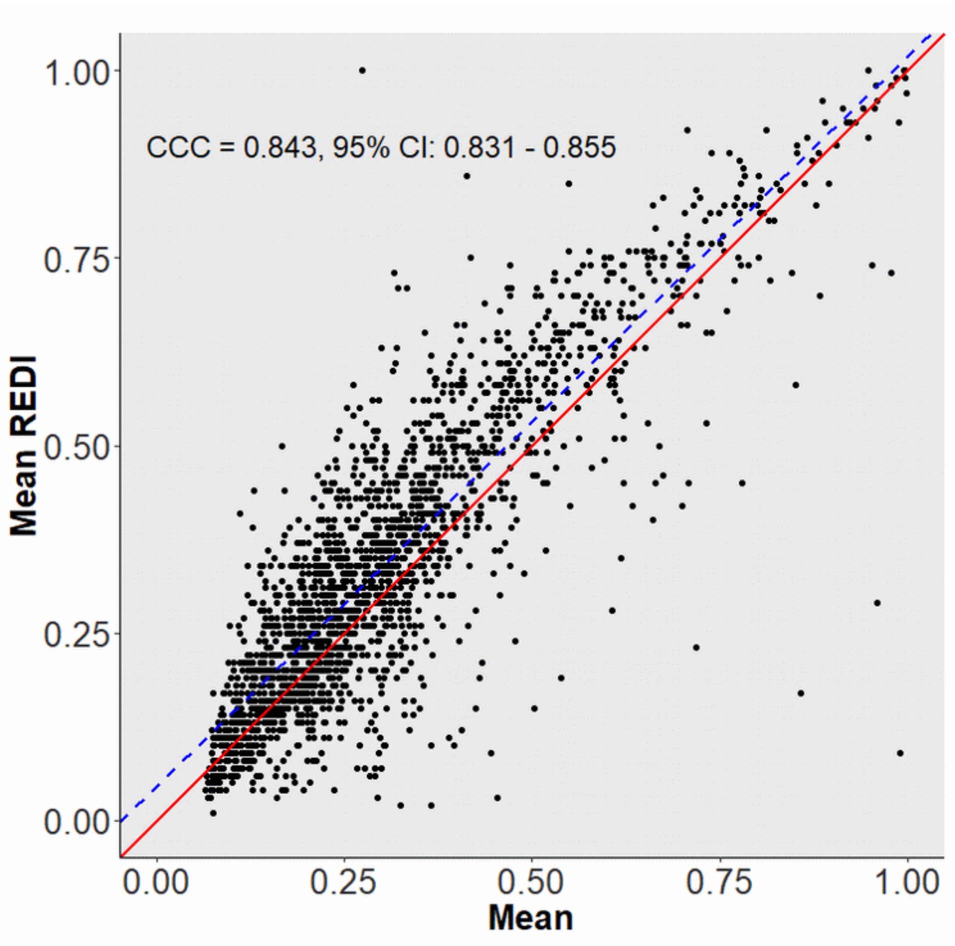

**Figure S4.** Overlap between CES and known miRNA binding sites from TargetScan
We found a total of 495 CES located within known miRNA binding sites from TargetScan v.7.2. The analysis was performed separately for broadly conserved, conserved and non conserved miRNA and miRNA binding site groups, as defined by TargetScan.

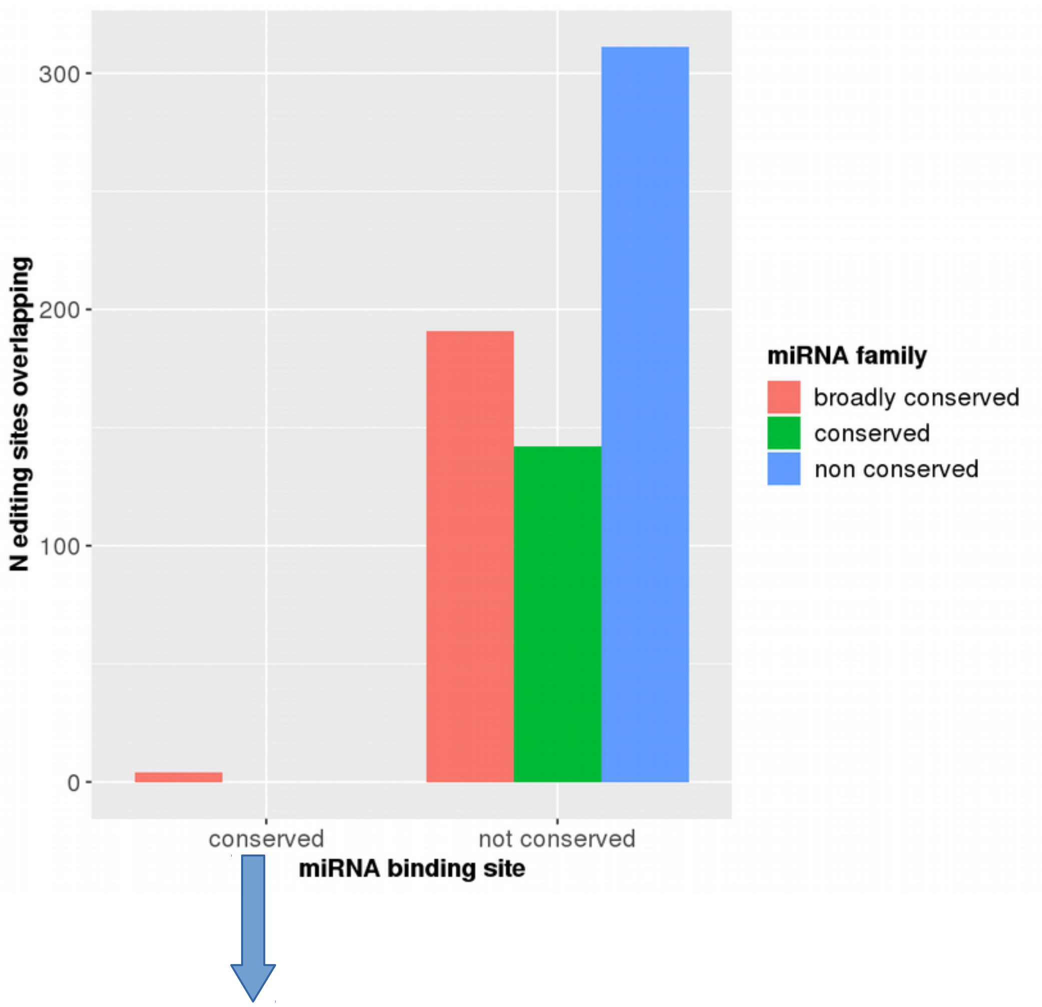

| | | |
|---|---|---|
| chr6 | 42905086 | CNPY3 |
| chr6 | 42905085 | CNPY3 |
| chr6 | 42905211 | CNPY3 |
| chr1 | 184762472 | FAM129A |

**Figure S5.** Correlation of editing levels between sites

Using Spearman rank test, we analyzed the correlation between editing levels among the 2,079 trusted sites. Significant correlations (FDR < 0.05) was generally low (a,d), except for close sites within 50 bp (b), for which we observed strong positive correlations (rho > 0.5). An example of editing island within 50bp on chromosome 3 is reported in correlation plot (c). Circle dimensions and color scale represent rho values of site-site correlation. Correlation and p-values for all relationship fir FDR < 0.05 are plotted in (d), with point colored according to distance between the sites considered. For the 66 couples of sites showing high correlations (rho > 0.5), the distributions of absolute difference in editing levels across subjects are reported in (e).

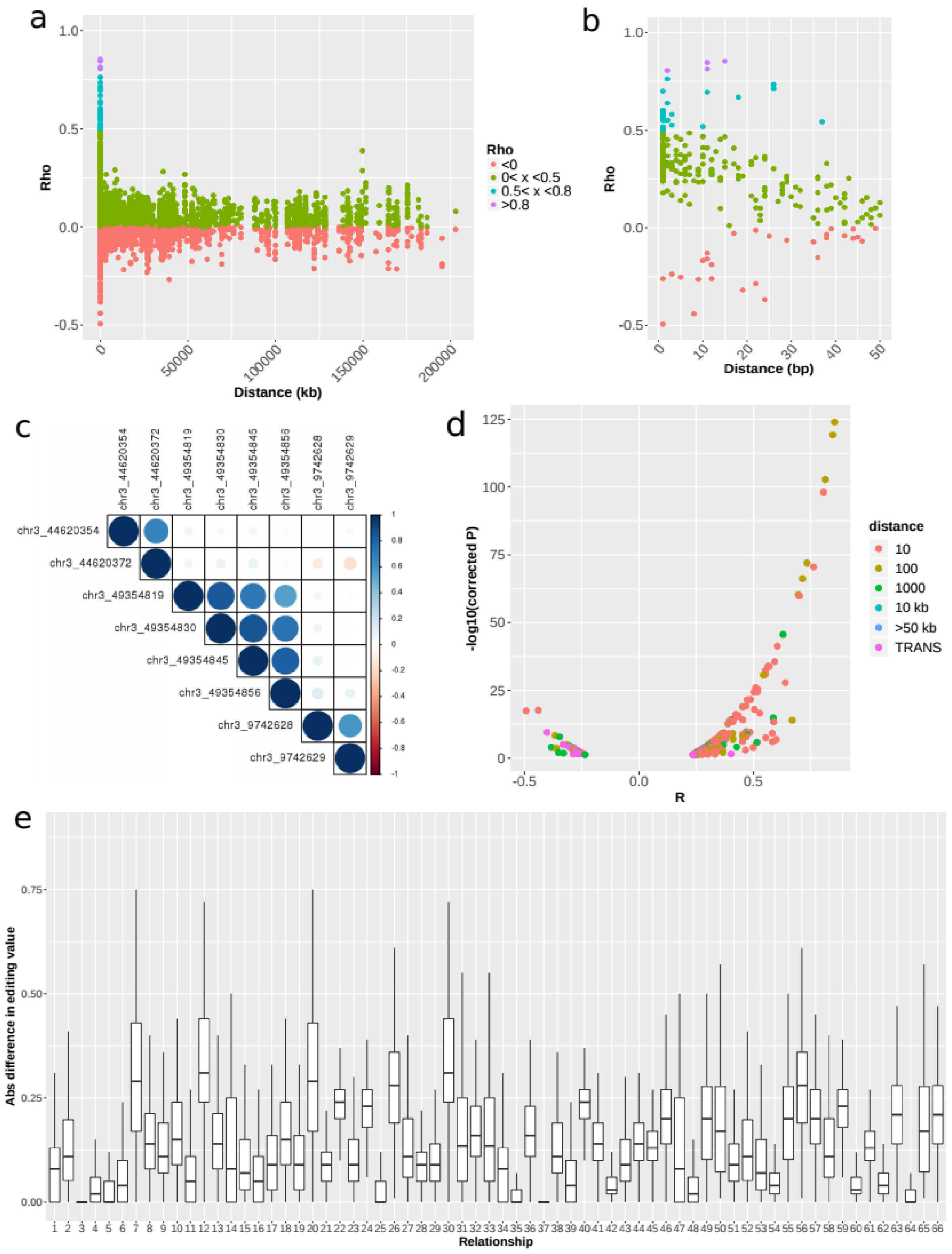

**Figure S6.** Correlation of *ADAR* and *ADARB1* expression on Alu and Non-Alu editing sites
We used robust regression analysis to estimate the association between *ADAR* (a) or *ADARB1* (b) expression and CES total editing rate, considering Alu and non-Alu sites separately. The graphs report adjusted p-value and $R^2$ value from robust regression analysis. *ADAR* showed association with both class of sites, while *ADARB1* showed no significant associations.

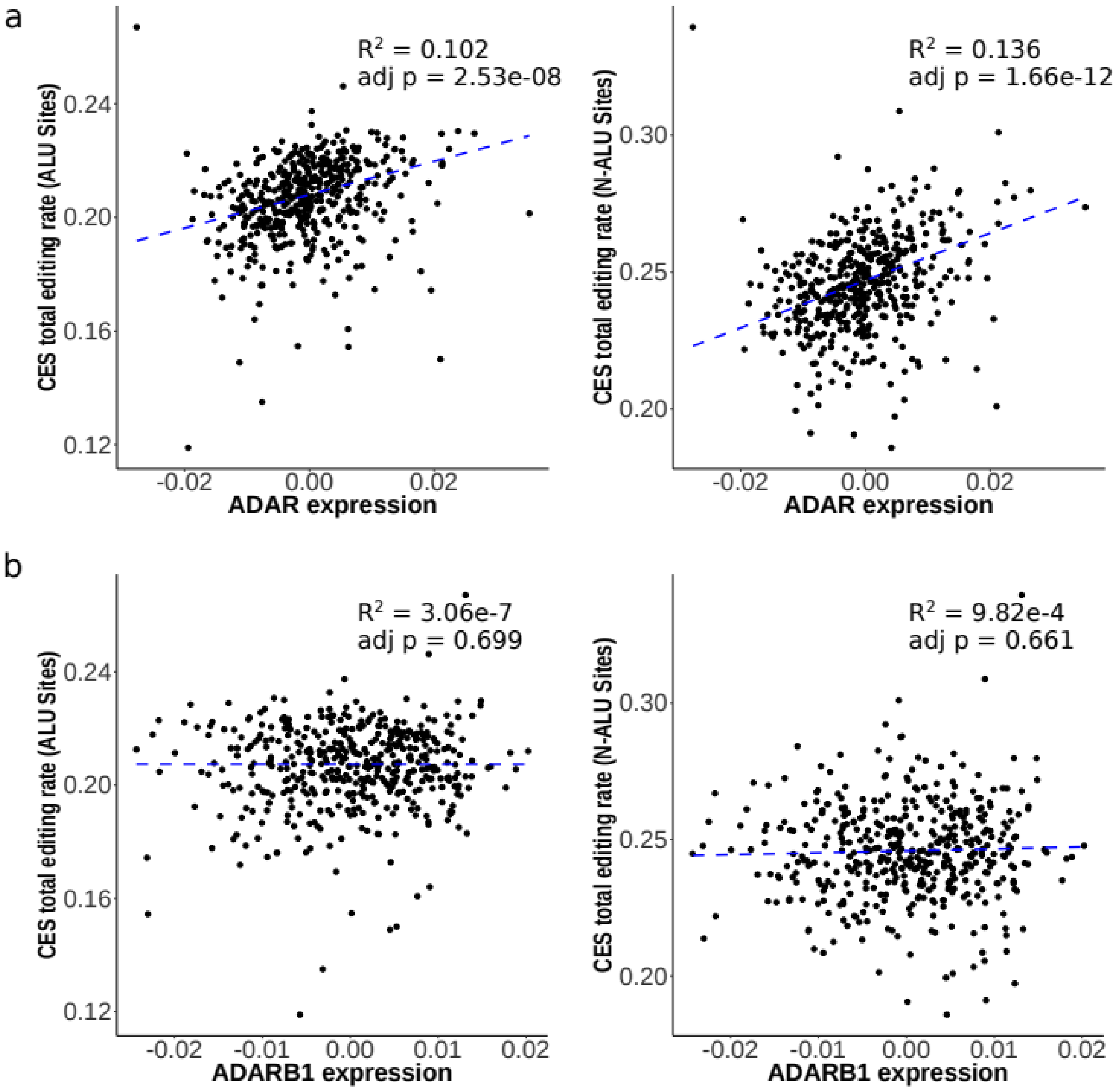

**Figure S7.** Linear regression models predicting *ADAR* / *ADARB1* expression level from cell composition variables and biological variables

To better evaluate the impact of cellular composition on *ADAR* / *ADARB1* expression, we computed a liner regression model predicting gene expression from the 7 cell variables (in red). We observed high correlation between predicted and expected values especially fro *ADARB1*. Adding the biological variables associated to *ADAR* / *ADARB1* expression (see Table S4) improve the prediction model for *ADAR*, but not *ADARB1* (in green).

R-squared values for each model are reported with the corresponding color. P-value refers to the LRT test between the model with and without the biological variables.

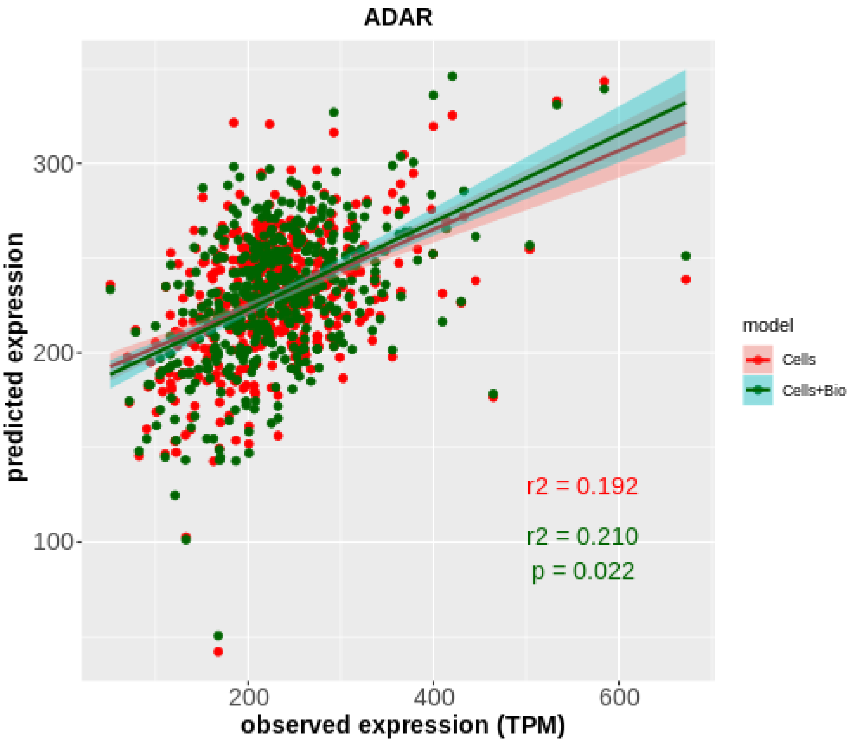

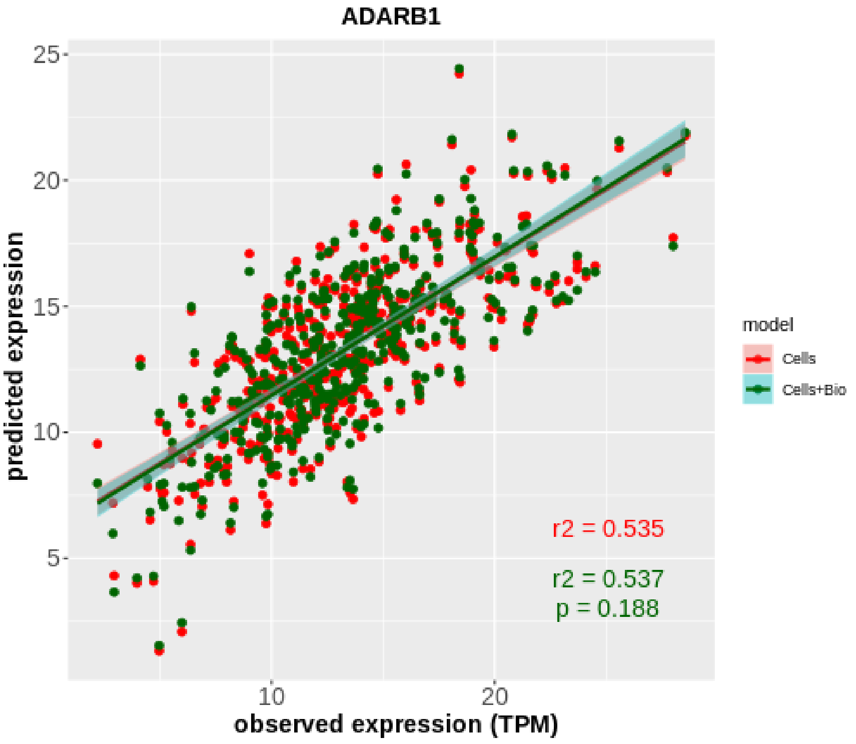

**Figure S8.** Effect of the top associated *ADAR* eQTL (rs6699825) on *ADAR* expression and CES total editing rate.

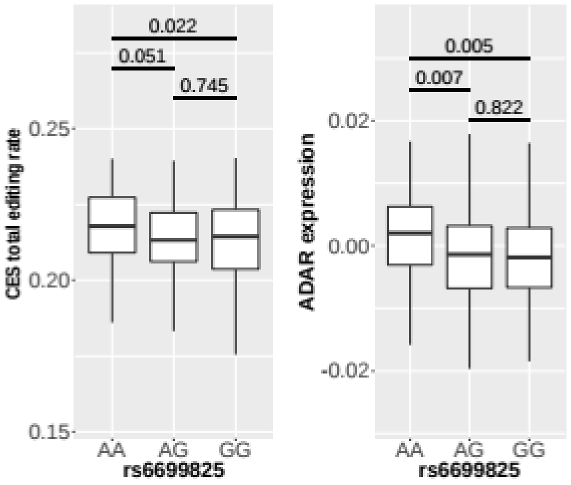

**Figure S9.** Distribution of the main RNA sequencing metrics for experiments used in this study.

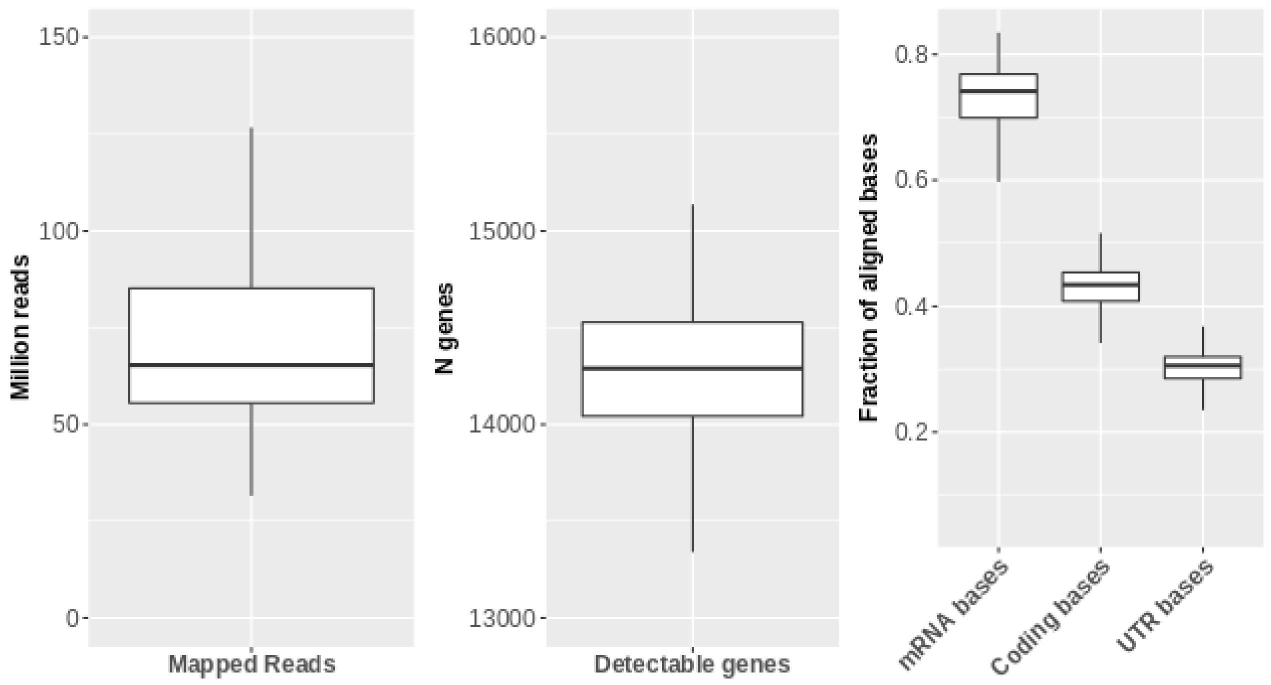

**Figure S10.** Correlation between observed editing variance and median coverage at editing sites. For single editing sites, we observed a strong correlation between median coverage and the observed variance, so that sites with lower coverage resulted in higher variance. The correlation seems to disappear for sites above 40 X coverage (blue dots). This suggest that site coverage should be taken into account when performing associations on editing levels.

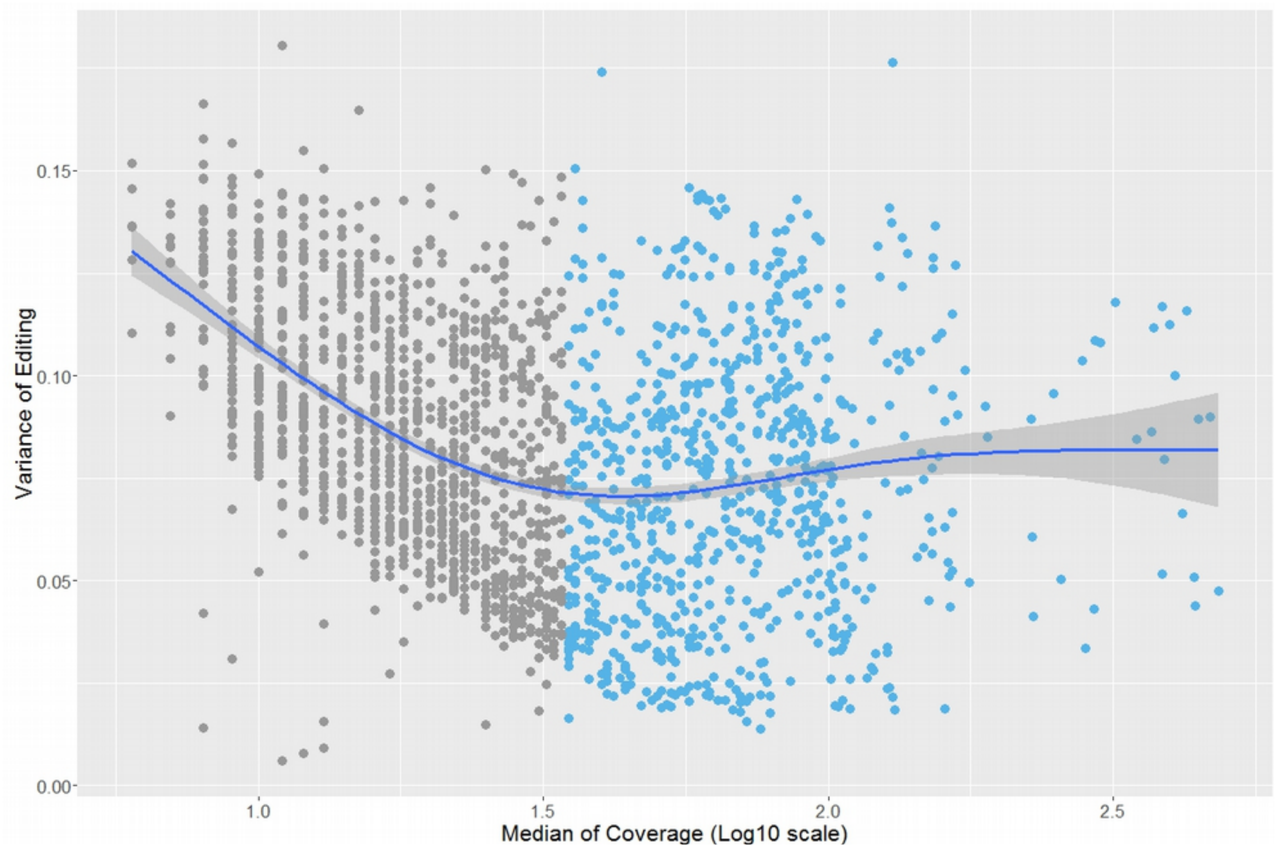
